# Mechanisms of SARS-CoV-2 Inactivation using UVC Laser Radiation

**DOI:** 10.1101/2023.02.03.526944

**Authors:** George Devitt, Peter B. Johnson, Niall Hanrahan, Simon I.R. Lane, Magdalena C. Vidale, Bhavwanti Sheth, Joel D. Allen, Maria V. Humbert, Cosma M. Spalluto, Rodolphe C. Hervé, Karl Staples, Jonathan J. West, Robert Forster, Nullin Divecha, Christopher J McCormick, Max Crispin, Nils Hempler, Graeme P. A. Malcolm, Sumeet Mahajan

## Abstract

Severe acute respiratory syndrome coronavirus 2 (SARS-Cov-2) has had a tremendous impact on humanity. Prevention of transmission by disinfection of surfaces and aerosols through a chemical-free method is highly desirable. Ultraviolet C (UVC) light is uniquely positioned to achieve inactivation of pathogens. We report the inactivation of SARS-CoV-2 virus by UVC radiation and explore its mechanisms. A dose of 50mJ/cm^2^ using a UVC laser at 266nm achieved an inactivation efficiency of 99.89%, whilst infectious virions were undetectable at 75mJ/cm^2^ indicating >99.99% inactivation. Infection by SARS-CoV-2 involves viral entry mediated by the spike glycoprotein (S), and viral reproduction, reliant on translation of its genome. We demonstrate that UVC radiation damages ribonucleic acid (RNA) and provide in-depth characterisation of UVC-induced damage of the S protein. We find that UVC severely impacts SARS-CoV-2 spike protein’s ability to bind human angiotensin-converting enzyme 2 (hACE2) and this correlates with loss of native protein conformation and aromatic amino acid integrity. This report has important implications for the design and development of rapid and effective disinfection systems against the SARS-CoV-2 virus and other pathogens.

## Introduction

The COVID-19 pandemic caused by the severe acute respiratory syndrome coronavirus-2 (SARS-CoV-2) (*1, 2*) spreads via nosocomial, public and work-place based infections (*3*). Transmission is thought to be direct via respiratory droplets or indirect via fomites and has led to increased interest in viral disinfection, including the use of ultraviolet (UV) light to inactivate virus in aerosols and on surfaces. The majority of studies into the effects of UV have focussed on the effects of UVA (400–320nm) and UVB (320– 280nm) due to the prevalence of these wavelengths in sunlight (*4*) and the availability of light sources in these ranges. More recently UVC (280–200nm) has seen increased interest as a method for viral inactivation in blood plasma extracts (*5, 6*). Short wavelength UVC (<240nm) shows increased efficacy for inactivation of MS2 phage (*7*), influenza (*8*), human coronaviruses (*9*) and damaging adenoviral proteins (*10*). These effects have been attributed to the absorption peak of proteins around 230nm (*11, 12*) but the effect of UVC on viral protein structure and function is not yet understood. This is important given the critical role that proteins play in viral cell entry (*13*).

Coronaviruses depend on a number of structural proteins for virus particle formation that include spike (S), membrane (M), envelope (E) and nucleocapsid (N) proteins. S forms trimeric projections from the surface of the virion, giving it its characteristic crown-like appearance. Each S subunit contains an S1 and S2 domain, mediating cell attachment and membrane fusion respectively. SARS-CoV-2 S has a receptor binding domain (RBD) within the C-terminal region of S1 that is responsible for binding human angiotensin-converting enzyme 2 (hACE2); an interaction driving SARS-CoV-2 cell tropism (*14*). Cleavage of the S, both at the S1–S2 boundary by furin, and within the S2 subunit by TMPRSS2 and cathepsin-L, primes the protein for membrane fusion (*15–18*). Owing to its dual role in mediating receptor binding and membrane fusion, we hypothesised S protein damage may be an effective route to viral inactivation.

Native conformation and structural integrity of proteins are required for correct function. The S protein-hACE2 binding involves a key sequence of amino-acid residues (*19–22*) constituting the receptor binding domain (RBD). The RBD conformation contains a twisted five-stranded antiparallel β sheet with short connecting helices (*23, 24*). Further, within this core, 2 α-helices and 2 β-strands, containing multiple key aromatic residues, and a disulphide bridge, form the receptor binding motif (RBM) (*21*). Aromatic residues (tryptophan, tyrosine and phenylalanine) absorb UVC light at 280, 275 and 258nm, respectively and disulphide bonds absorb at 260nm (*25*). Therefore the RBM and the spike protein as a whole, along with other viral proteins, is potential target for UVC induced damage.

In this work we demonstrate UVC inactivation of SARS-CoV-2 virus and its dose dependence. To understand the mechanism of UVC inactivation we use two UVC wavelengths (266nm, near UVC; 227nm, far UVC) from solid-state continuous wave lasers. 266nm radiation is strongly absorbed by nucleic acids (*26*) and by the aromatic rings of amino acid sidechains while 227nm is absorbed less than 266nm by nucleic acids, but more strongly by proteins (*27*). We explore the dose dependent damage to viral RNA and its scaling with genome size. We probe in-depth the effect of UVC on the SARS-CoV-2 S protein using binding assays, morphological characterisation and molecular spectroscopy. UVC irradiation of SARS-CoV-2 S protein severely affected its ability to bind to hACE2 which correlated well to the observed changes in protein conformation and oxidation of aromatic residues. We postulate that the UVC induced damage to proteins, in combination with that to RNA, is an important contributor towards the inactivation of the SARS-CoV-2 virus with immediate implications for wavelength and dose selection for preventing transmission of Covid-19 and other airborne pathogens.

## Results and Discussion

### SARS-CoV-2 inactivation using 266nm laser

To examine the impact of UVC on SARS-CoV-2 infectivity, virus was dried on to polystyrene surfaces which were then exposed to 266nm irradiation at 0.5mW/cm^2^ for between 1 and 100s. The exposed virus samples were resuspended and assessed by plaque assay. A clear dose dependent response was seen, with the highest dose (50mJ/cm^2^) leading to 99.89% inactivation (**Figure 1A, B** and **Supplementary S1**). In a separate experiment ^a^ following a 25mJ/cm^2^ (1s, 25mW/cm^2^) dose, only a single infectious virion across 3 technical repeats was recovered, equating to 99.97% inactivation. Doses of 75mJ/cm^2^ (25mW, 3s) or greater reduced infectivity below the limit of detection of 0.01% (**Supplementary S2**). Thus surface decontamination of SARS-CoV-2 by 266nm UVC can be achieved within seconds at modest UVC doses.

**Figure 1.**
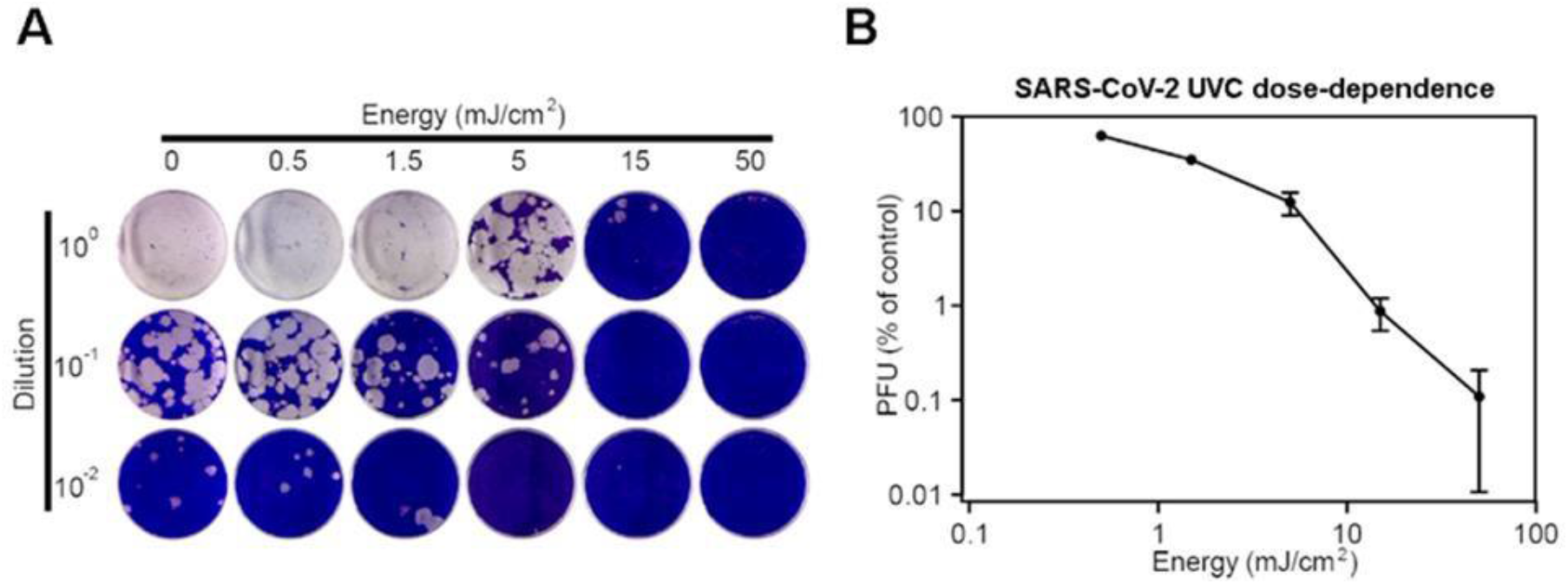
SARS-CoV-2 virus surface inactivation by 266nm UVC continuous wave laser. A) Representitative set of images from a plaque assay showing the dose (horizontal) and serial dilution (vertical). B) Dose-dependence of SARS-CoV-2 virus to 266nm UVC radiation, from two independent experiments performed in triplicate.

For comparison the UVC inactivation dose observed is higher than the ≤2mJ/cm^2^ reported for human coronaviruses alpha HCoV-229E, beta HCoV-OC43 (*8*) and H1N1 influenza (*9*). This could be due to the use of 222nm radiation in those studies, or due to a much shorter path length (≤1μm aerosols vs. dried 25μL media droplets used here). Additionally, the above coronaviruses use human aminopeptidase N (*28*) and sialic acid sugars (*29*) as a receptor for cell entry, whilst SARS-CoV-2 virus spike protein is optimised for binding hACE2 (*30*). Viral cell entry, mediated by the spike protein, and replication, facilitated by the genome, are both essential parts of the infection processes (**Figure 2A**). Therefore, we explored the effect of UVC on macro-molecular components involved in SARS-CoV-2 viral infection.

**Figure 2.**
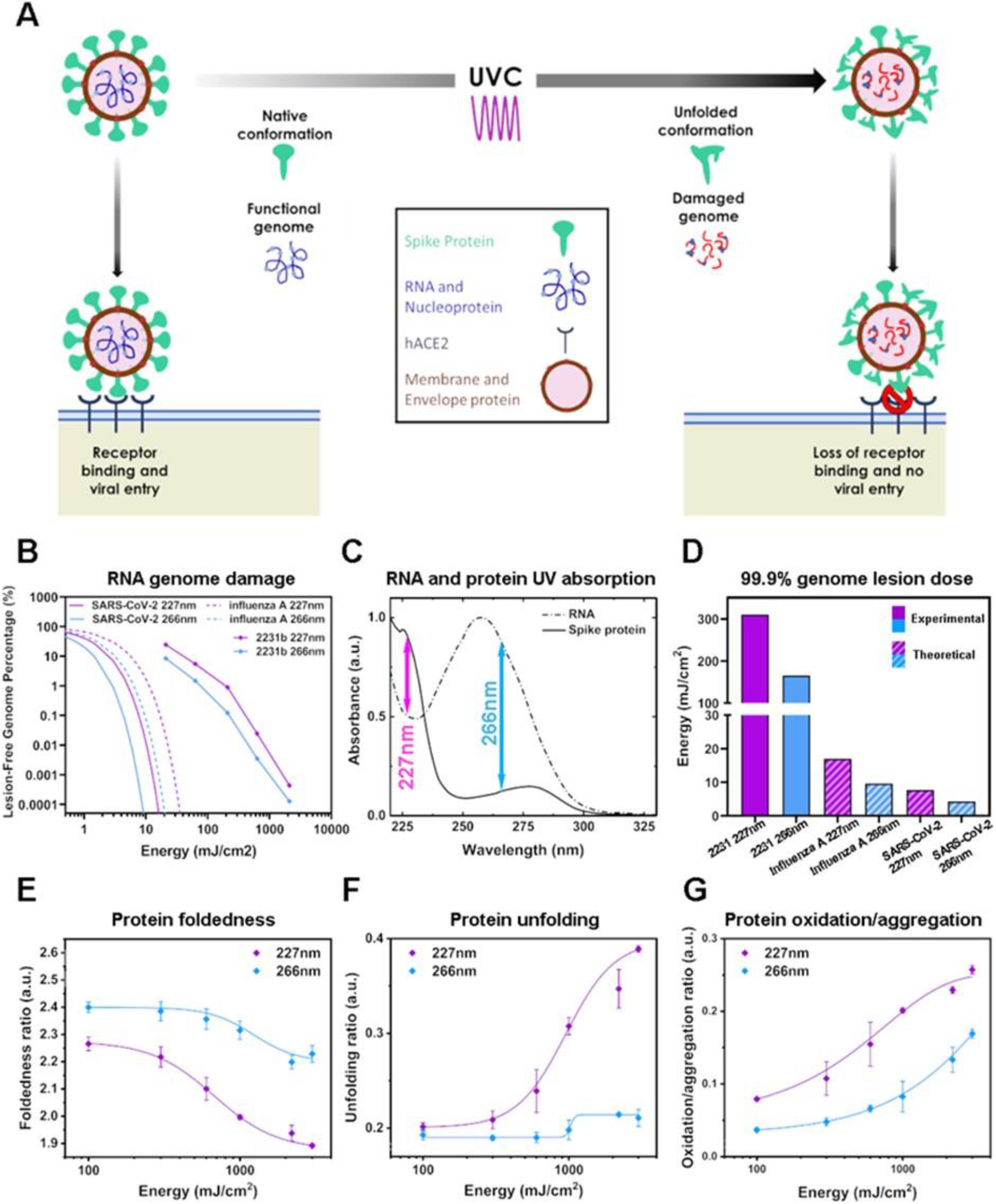
UVC dose dependence of ssRNA and rSARS-CoV-2 S protein integrity. A) Schematic depicting the effect of UVC radiation on SARS-CoV-2 cellular entry and reproduction. B) Dose-dependent RNA inactivation. Figure depicts RT-qPCR product off ߢkb RNA template, and projected 30kb RNA template as a percentage of non-irradiated RNA control. n=3, plotted data represents mean and SD from 6 cycle threshold readings. C) UV absorption spectrum of rSARS-CoV-2 S. Arrows indicate UVC laser wavelengths used in this study. D) Experimentally determined 227nm and 266nm UV dose requirements for 99.9% replication inhibition from the 2,231 base MS2 amplicon, and theoretical model extrapolated using Poisson statistics for larger 13,588 and ~30 kb SARS-CoV-2 genomes. E.) Dose-dependance of rSARS-CoV-2 S foldedness ratio to UVC radiation determined by UV-vis absorption spectroscopy. Foldedness ratio = A280/A275 nm + A280/A258 nm. F) Dose-dependence of rSARS-CoV-2 S unfolding ratio to UVC radiation determined by UV-vis absorption spectroscopy. Unfolding ratio = A280/A230 nm. G) Dose-dependance of SARS-CoV-2 S oxidation/aggregation ratio to UVC radiation determined by UV-vis absorption spectroscopy. Oxidation/aggregation ratio = A320/A280 nm. n=2, the plotted data represent the average and SD from 10 absorbance readings for native rSARS-CoV-2 Sand 3–4 absorbance readings per UVC condition.

### UVC damage of RNA

227nm and 266nm UVC wavelengths were used to test the effect on RNA integrity using the MS2 ssRNA viral genome (3.57 kb) and an RT-qPCR assay (*7*). Here we define RNA damage as that sufficient to prevent *in vitro* reverse transcriptase progression, however, we note that host ribosomal sensitivity to damaged RNA may differ. Damage therefore results in less cDNA template for the subsequent PCR reaction. Our assay could detect RNA at concentrations up to 6 orders of magnitude below the stock concentration. We found that both wavelengths damaged RNA, in a dose dependant manner (**Figure 2B**, 99.9% reduction dose: 227nm, 285mJ/cm^2^; 266nm, 180mJ/cm^2^), but 266nm was more effective, presumably as a result of the strong absorption by RNA around 260nm (**Figure 2C**).

In our assay the PCR primers amplified targets 925 bases and 2231 bases distal from the primer used for first strand synthesis. Thus, our assay probed RNA integrity only for these lengths, whilst the SARS-CoV-2 genome is ∼30kb in length. Using a probabilistic interpretation involving the Poisson statistic that considers that UV-mediated RNA lesion events are proportional to genome length we modelled our results to predict the required dose for inactivating the SARS-CoV-2 genome (**Figure 2B, D, Supplementary Figure S3)**. To damage 99.9% of SARS-CoV-2 genome the model predicts a dose of only 7.7mJ/cm^2^ at 227nm or 4.25mJ/cm^2^ at 266nm. These are not dissimilar to the dose needed to inactivate other human coronaviruses using similar wavelengths (*9*). However, the viral inactivation dose observed for SARS-CoV-2 was ∼11.7 times higher (∼50mJ/cm^2^ at 266nm, **Figure 1B**). Proteins also absorb UVC radiation in this spectral range (**Figure 2C**). Hence, we next investigated the possibility that UVC induced damage to proteins in the virus particles, with implications for structural integrity and capacity for cell entry.

### UVC damage of SARS-CoV-2 spike protein

We expressed solubilized and stabilized recombinant SARS-CoV-2 spike protein (rSARS-CoV-2 S). The rSARS-CoV-2 S variant replaces the furin cleavage site with a “GSAS” linker and, along with several other mutations, helps to stabilize the trimeric structure of the spike protein (*24*). The mutations are required to maintain native-like structure and we note that this increased stabilisation likely sets a higher threshold for UVC-mediated inactivation. rSARS-CoV-2 S has two peaks in its UV-vis absorption spectrum, which can be targeted by the two wavelengths used in this study (**Figure 2C**). We established a UVC dose-response for each wavelength for rSARS-CoV-2 S protein integrity using UV-visible absorption spectra as a readout (**Figure 2E-G**). The conformational state of a protein affects its absorption properties, thus protein unfolding, oxidation and aggregation are all reflected by changes in the absorption spectrum (*31*). We observed a dose-dependent decrease in the foldedness ratio of rSARS-CoV-2 S (A280nm/A275nm + A280nm/A258nm) (*25*) with increasing dose of UVC radiation, with 227nm producing a greater dose matched effect than 266nm (**Figure 2E**). The A230nm peak is sensitive to protein secondary structure, with an increase in intensity observed upon protein unfolding (*31*). We observed a large dose-dependent increase in the A230nm/A280nm ratio for rSARS-CoV-2 S in response to 227nm, with a subtle increase in response to 266nm irradiation (**Figure 2F**). This suggests that 227nm irradiation damages secondary structure more than 266nm irradiation. Similarly, the A320nm/A280nm ratio increases upon protein aggregation (*32*) or upon oxidation of tryptophan to n-formylkynurenine (NFK) (*33*). We detected a UVC dose-dependent increase in A320nm/A280nm ratio (**Figure 2G**), again consistent with loss of native protein conformation. Taken together S protein unfolding and conformational loss due to UVC exposure are predicted to impair functionality.

### UVC induced loss of rSARS-CoV-2-hACE2 binding

In order to correlate the changes caused by UVC treatment to the functionality of the rSARS-CoV-2 spike protein we selected low (100mJ), medium (600mJ) and high (2200mJ) UVC doses (**Supplementary Figure S4**) and performed surface plasmon resonance (SPR) binding assays. To determine the ability of rSARS-CoV-2 S to bind hACE2, both with and without UVC treatment, the protein was expressed with a C-terminal HisTag which allows for both purification and also binding to a nickel-coated SPR sensor surface.

Serial dilutions of hACE2 were sequentially flowed over the S protein-coated chip and the response measured (**Figure 3A and 3B**). Untreated rSARS-CoV-2 S protein bound hACE2 with a K_D_ of 54nM (**Figure 3C and Supplementary Table 1**). Using a flow rate of 10μl/min for 240s resulted in a calculated R_max_ of 377.4. This value reflects the amount of rSARS-CoV-2 S protein which has successfully bound to the chip. Both UVC wavelengths, 227nm and 266nm, resulted in an increase in K_D_ and a concurrent decrease in R_max_ (**Figure 3D-3I**) indicating higher dissociation and reduced binding. At the highest dose tested (2200mJ/cm^2^) the R_max_ was only 2% compared to that of untreated rSARS-CoV-2 S protein for 227nm and 6% for 266nm irradiation which are similar to the negative control. This is likely due to degradation of the S protein, inhibiting hACE2 binding. The 600mJ dose resulted in both an increase in K_D_, reflecting a decrease in affinity of rSARS-CoV-2 S protein for hACE2, and a R_max_ decrease, with the 227nm wavelength being more effective than the 266nm wavelength. The 100mJ dose resulted in a smaller increase in K_D_ and decrease in R_max_ compared to the 600mJ dose, but at this dosage both 227nm and 266nm wavelengths performed similarly. While UVC dose dependent reduction in binding is clear, the largest percentage decrease in binding capacity occurs within 100mJ itself (**Figure 3B**). The higher dose required to inactivate the recombinant SARS-CoV-2 spike protein in comparison to the virus may be due to the synergistic effect of UVC induced RNA damage, as well as differences in stability of the recombinant and wild-type viral spike protein. That UVC damages the spike protein, reduces its binding ability and thus will diminish viral entry is clear. Therefore, UVC wavelengths, which damage both viral entry and replication, are likely to be very powerful disinfectants for viruses such as SARS-CoV-2.

**Figure 3.**
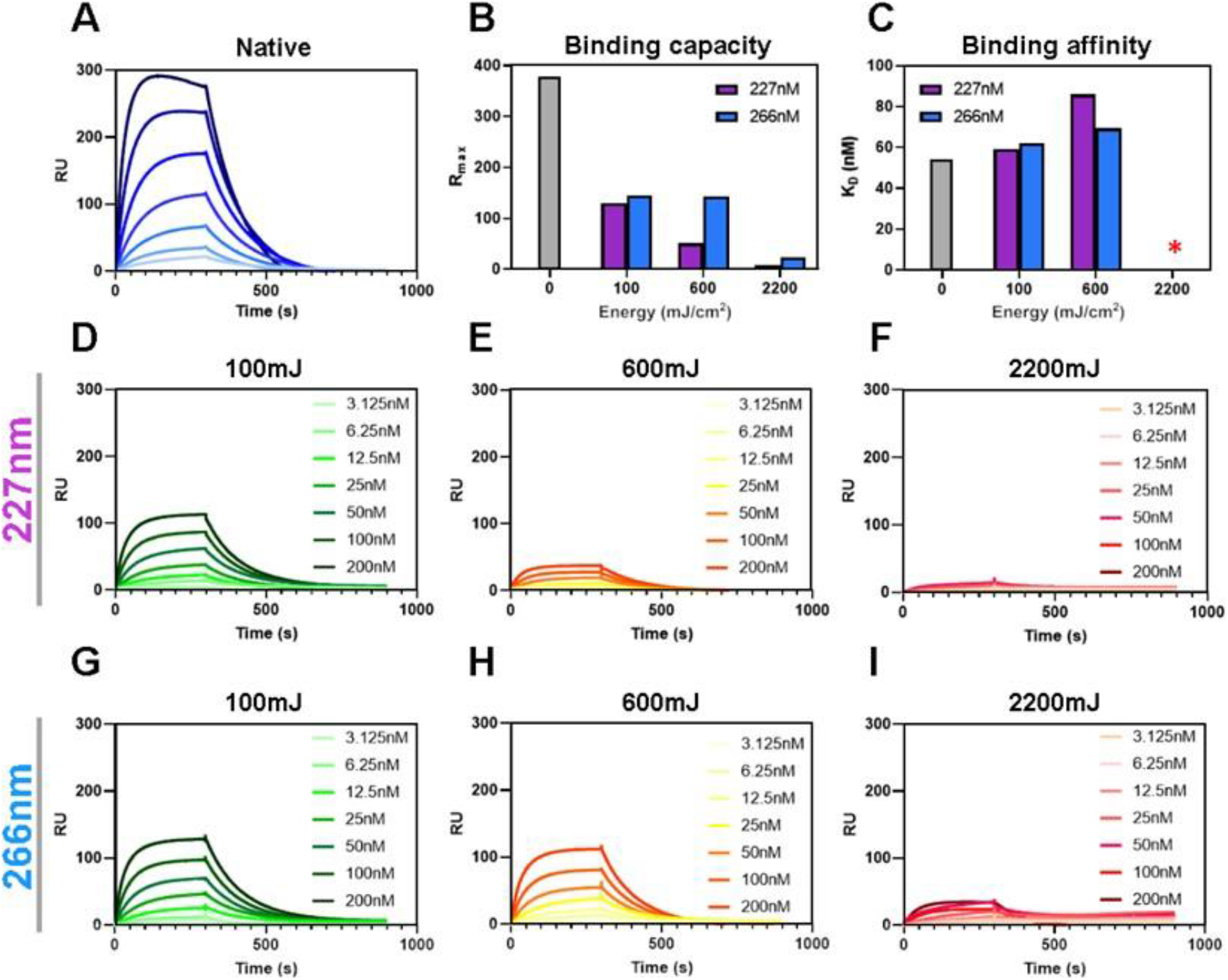
Surface plasmon resonance (SPR) analysis of rSARS-CoV-2 trimeric spike glycoprotein shows the impacts of UVC treatment on hACE2 binding. A) Sensogram depicting the binding of solubilized recombinant SARS-CoV-2 to serial dilutions of the hACE2 receptor with dilutions ranging from 200nM to 3.125nM. Data represent an average of three analytical repeats. B) Comparison of the calculated Rmax values for each UVC treatment condition expressed as percentage of the non-irradiated protein. C) K¬D values for all of the SPR experiments performed. This parameter was calculated using a 1:1 binding model using Biacore Evaluation software. The response for the 2200mJ dose for 227nm and 266nm was not sufficient to accurately determine KD. D-I) SPR experiments were performed identically to panel A except an equal amount of S protein was treated with the dosage indicated.

### UVC radiation has distinct effects on rSARS-CoV-2 S conformation

In order to understand the reduction of rSARS-CoV-2 S function, we investigated the global changes in protein conformation induced by UVC. We first describe the effect with 227nm and then 266nm.

Hydrophobic residues buried within the native protein structure can be exposed upon protein unfolding. Hence, to measure surface hydrophobicity we used a Bis-ANS (4,4’-dianilino-1,1’-binaphthyl-5,5’-disulfonic acid dipotassium salt) binding assay. Increased Bis-ANS binding and fluorescence was observed with 227nm UVC irradiation (**Figure 4A**), however, this increase in hydrophobicity became smaller with increasing doses. This may be because Bis-ANS has lower affinity for oxidised proteins (*34*). Therefore, we probed rSARS-CoV-2 S for oxidation sensitive tryptophan residues (reduced tryptophan fluoresces at 330nm and its oxidised product, N-formyl kynurenine (NFK), fluoresces at 425nm). We observed a dose-dependent decrease in the 330nm/425nm emission ratio (**Figure 4B**). This decreasing ratio is primarily due to a loss in tryptophan fluorescence, with a large increase in NFK fluorescence only occurring at the highest dose (**Figure 4C**).

**Figure 4.**
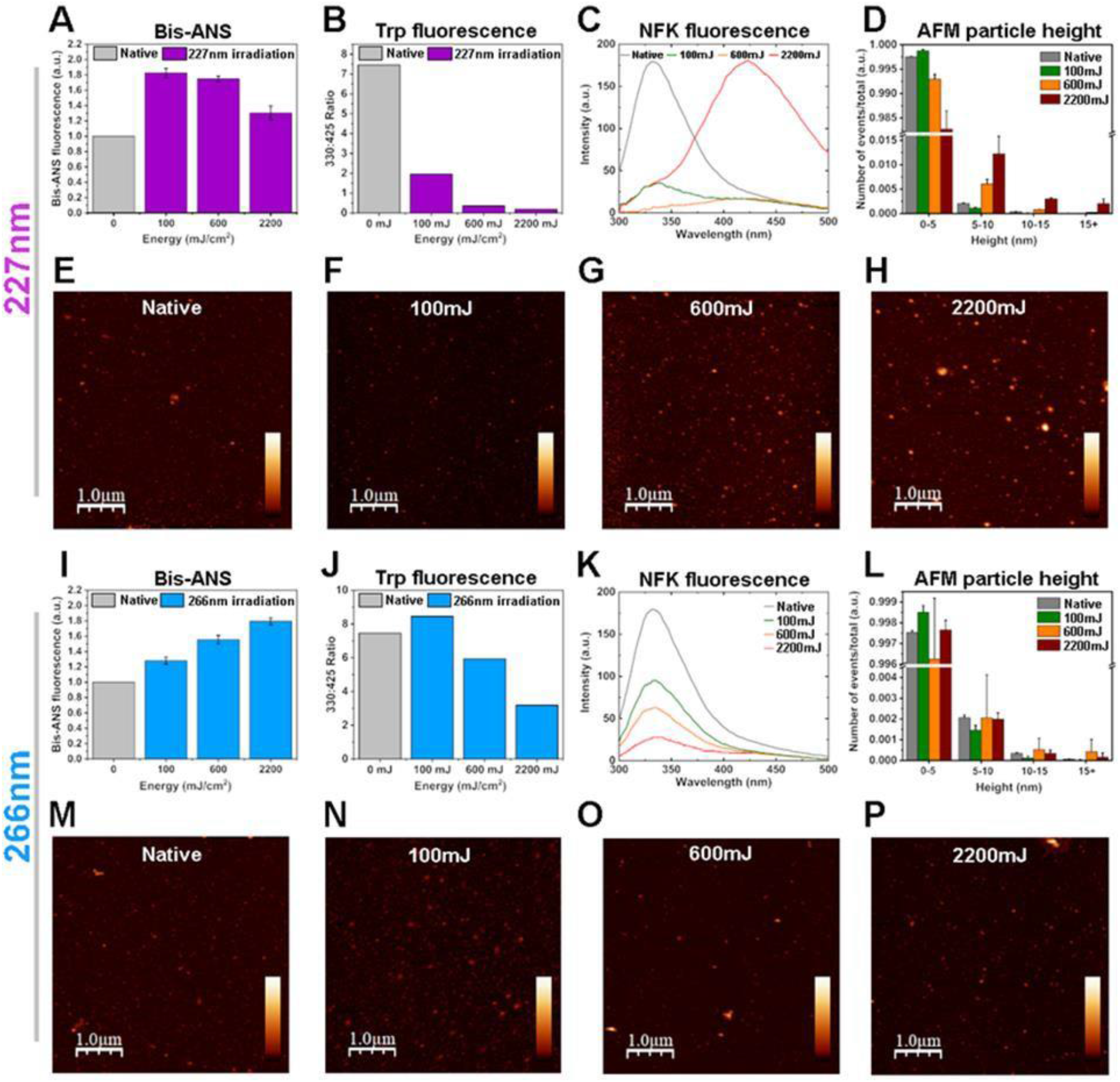
UVC dose-dependent loss of rSARS-CoV-2 spike protein conformation. Readouts of rSARS-CoV-2 spike protein conformation induced by 227nm (A-H) or 266nm (I-P) radiation A,I) Bis-ANS binding measured by fluorescence emission at 490nm. n=2, the plotted data represent the average and SD from 4 fluorescence readings. B,J) Fluorescence emission ratio 330/425nm representing tryptophan/NFK fluorescence. C,K) Fluorescence emission spectra from 300-500nm representing changes in intrinsic tryptophan fluorescence. n=1, plotted data represents the average from 3 fluorescence readings. D,L) Particle height analysis of atomic force microscopy (AFM) images. 3-4 images were used for each condition. n=2, the plotted data represent the average and SD from 2-4 AFM images. E-H, M-P) Representative tapping-mode AFM images of SARS-CoV-2 spike protein bound to a mica surface after irradiation with 0mJ (E,M), 100mJ (F,N), 600mJ (G,O), 2200mJ (H,P). Images are 5x5μm, scale bars represent 1μm and z-scale is equal to 0-30nm.

Protein unfolding and the exposure of hydrophobic residues can lead to protein aggregation (*35*). We therefore used atomic force microscopy (AFM) to probe the protein morphology. Particle height analysis was performed on AFM images to quantify the number of protein aggregates induced by 227nm irradiation (**Figure 4D**). Medium and high doses of 227nm irradiation induced protein aggregation, with a transition from smaller structures (<5nm) to larger assemblies (5-50nm). The morphology of these assemblies can be observed in the representative AFM images, which show an increase in amorphous aggregates after exposure to medium and high doses of 227nm radiation (**Figures 4E-H, Supplementary S5**).

Interestingly, higher doses of 266nm radiation did not induce the same effects as 227nm radiation on protein conformation. We observed a dose-dependent increase in Bis-ANS binding (**Figure 4I**), in contrast to 227nm radiation. The same trends were observed using bovine serum albumin (BSA) irradiated by 227nm and 266nm wavelengths (**Supplementary S6**). In order to assess whether this difference was due to the lower efficiency of the 266nm radiation on damaging protein conformation, we irradiated rSARS-CoV-2 S with a dose of 10J (∼4.5 fold greater than the high dose), which resulted in a further increase in Bis-ANS binding (**Supplementary S7**). In line with this, the 330nm/425nm ratio from tryptophan fluorescence showed a lesser decrease after 266nm irradiation in comparison to 227nm irradiation (**Figure 4J**), with no visible NFK fluorescence being observed at 425nm (**Figure 4K**). Further, 266nm radiation does not cause the same amount of amorphous protein aggregation as 227nm radiation at the doses used (**Figure 4L-P**). Together this shows that both wavelengths can cause protein damage and unfolding, however 227nm irradiation of SARS-CoV-2 causes distinctly higher tryptophan oxidation and amorphous protein aggregation compared to 266nm irradiation, which correlates with a greater reduction in hACE-2 binding.

### 227nm UVC damages rSARS-CoV-2 secondary and tertiary structure

To determine the specific molecular mechanisms of UVC-induced unfolding of rSARS-CoV-2 S for 227nm and 266nm radiation, we probed for vibrational changes using Raman spectroscopy. The Raman spectrum includes several regions sensitive to protein conformation: Amide I at ∼1650cm^-1^, an established marker of secondary structure; disulphide bonding at ∼500cm^-1^; and aromatic ring vibrations throughout the Raman spectrum, sensitive to tertiary structures (*36*).

We acquired Raman spectra of the untreated and UVC treated proteins and analysed the above-mentioned regions of the spectrum (**Supplementary S8**). The second derivative of the Amide I spectra allows subtle changes to be highlighted and ‘hidden peaks’ revealed (*37*). The second derivative spectrum for native rSARS-CoV-2 S (**Figure 5A**, grey trace) shows strong peaks at 1650cm^-1^ (α-helix), 1672cm^-1^(β-sheet) and a small peak at 1693cm^-1^ (turns). This is in line with the crystal structure of SARS-CoV-2 S (*24*). Upon irradiation with 227nm wavelength, rSARS-CoV-2 S shows a dose dependant loss in α-helix and β-sheet vibrations, with a concurrent increase in intensity ∼1685cm^-1^, corresponding to nonregular structure. 227nm (100mJ) causes loss of turn structures. 227nm UVC irradiation also causes a dose-dependent decrease in the ∼509cm^-1^ peak demonstrating a loss of disulphide bonds (**Figure 5B**). Thus given that the SARS-CoV-2 RBM contains two β-strands, 2 α-helices and a disulphide bond (*21*), these data correlate well with the observed loss in hACE-2 binding (**Figure 3**).

**Figure 5.**
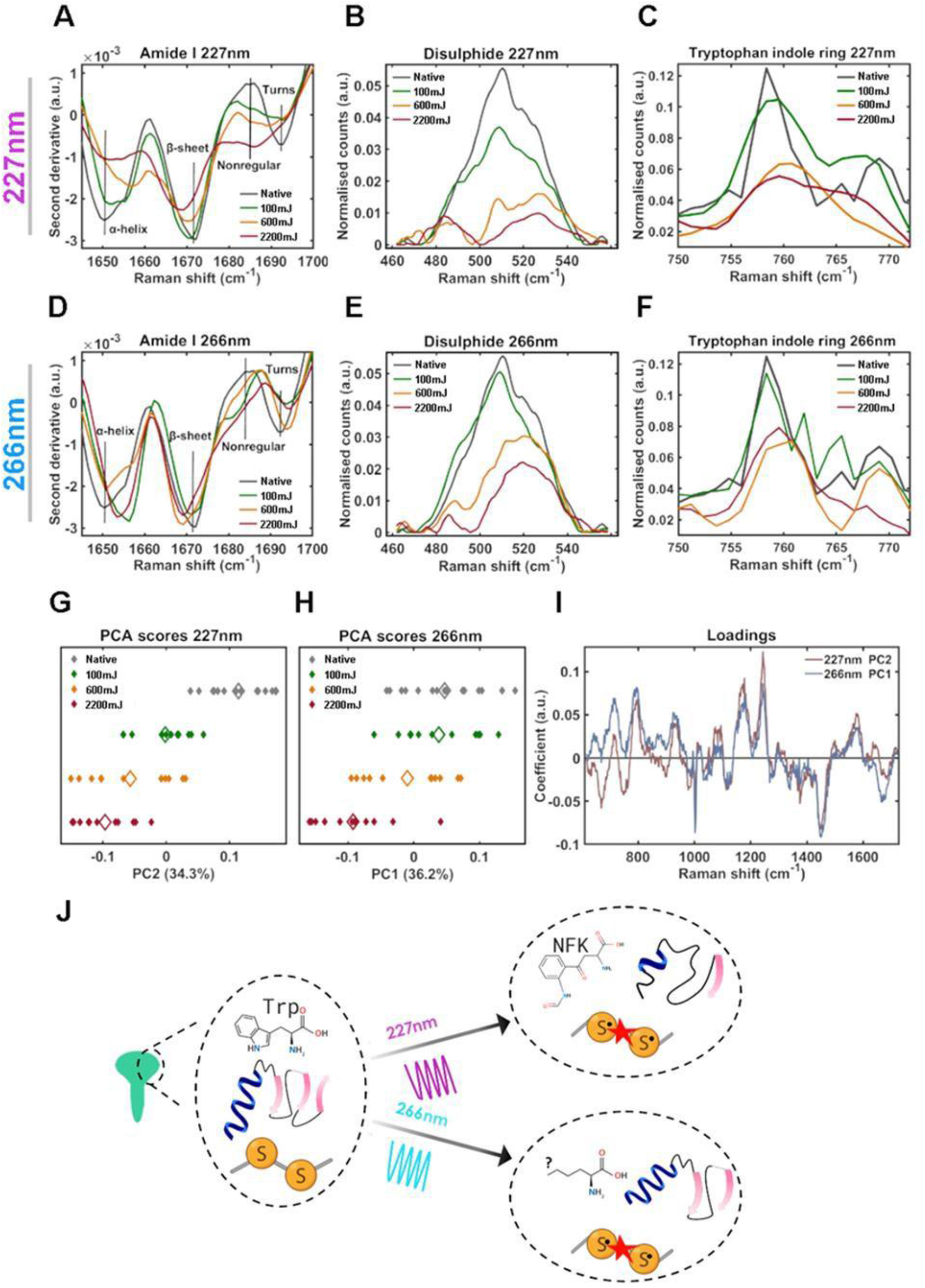
Mechanism of UVC-induced rSARS-CoV-2 S conformational damage. Raman spectra of rSARS-CoV-2 S protein following irradiation by 227nm (A-C) or 266nm (D-F) light. A,D) Amide I second derivative spectra from 1645-1700cm-1 B,E) Disulphide region spectra from 462-558cm-1. C,F) Tryptophan indole ring spectra from 750-772cm-1. G-H) 1-dimensional principle component analysis (PCA) scores plot of Raman spectra for 227nm (G) and 266nm (H) irradiated rSARS-CoV-2 S. Each solid diamond represents the PC score of a single spectrum. Hollow diamonds represent mean score. I) PC loadings spectra representing the spectral variation responsible for the score across the given PC axis. J) Schematic depicting conformational damage induced by 227nm and 266nm radiation determined by Raman spectroscopy. n=2, plotted spectra represent the class means of 10-15 spectra per class.

There are a number of aromatic amino acid markers in the Raman spectrum, including those for phenylalanine, tyrosine and tryptophan. We observe a dose dependent decrease in the tryptophan indole ring vibration 760cm^-1^ peak with increasing 227nm dose, as well as peak broadening, suggesting a loss in the ordered ring structure of tryptophan (**Figure 5C**) This suggests tryptophan oxidation (*38*) and protein denaturation (*39*). Further evidence of protein denaturation is suggested by the dose-dependent increase in the ratio of 830cm^-1^/850cm^-1^ (**Supplementary S9**). This corresponds to the Fermi resonance vibration in tyrosine and indicates changes in the tyrosine hydrogen bonding environment (*40*) that correlate well with the Bis-ANS data (**Figure 4**). Together, this suggests that irradiation of rSARS-CoV-2 S by 227nm light causes a loss in secondary and tertiary structure, resulting in denaturation of the protein and is consistent with its loss of function.

### 266nm UVC primarily damages SARS-CoV-2 tertiary structure

We similarly assessed conformational changes induced by 266nm irradiation in the spike protein. While a dose-dependent loss in α-helix and β-sheet peak intensity was not observed (**Figure 5D**) an overall loss in the peak intensity of turns and an increase in nonregular structure was observed. Thus, unfolding occurs but to a lesser degree than with 227nm irradiation, which is in agreement with UV-vis absorption results (**Figure 2F**). A dose-dependent decrease of the disulphide peak (**Figure 5E**, 509cm^-1^), as well as in the tryptophan indole ring vibration peak (**Figure 5F**, ∼760cm^-1^) were observed. Together, this suggests that irradiation of rSARS-CoV-2 S by 266nm light causes a loss in native conformation, although secondary structural components are less effected when compared to 227nm radiation. Disulphide bonds absorb weakly at ∼260nm (*41*) but excitation energy transfer (EET) (*42*) from tryptophan and tyrosine can cause their cleavage (*43, 44*). As both 227nm and 266nm radiation cause disulphide breakage in rSARS-CoV-2 S, it is likely that the cystine bonds are being targeted indirectly by EET due to their vicinity to Trp and Tyr residues.

### 227nm and 266nm UVC degrade aromatic rings

S protein tyrosine residue (Y484) is centrally involved in hACE-2 binding (*45*). Hence, we assessed the effect of UVC radiation on pure preparations of tyrosine, as well as tryptophan. Both 227nm and 266nm radiation yielded a dose-dependent loss of aromatic ring conformation (**Supplementary S10**), broadening of UV-absorption peaks, as well as the loss of sharp aromatic vibration peaks in the Raman spectrum, which were replaced by broad C-C bond vibrations ∼930cm^-1^ (**Supplementary S9B**). Together, this suggests, as expected, that UVC irradiation causes degradation of the aromatic rings in Tyr and Trp (*46*).

### 227nm and 266nm-induced protein unfolding have similar molecular mechanisms

In order to identify, in an unbiased manner, differences in mechanism between 227nm and 266nm radiation we performed principal component analysis (PCA) on the Raman spectra and selected components representing UVC dose-dependent changes (**Figure 5G, 5H**). Interestingly, both loadings spectra show that the same features, albeit with differing coefficients, are responsible for UVC dose-dependent changes (**Figure 5I**). The same result was observed for PCA of the normalised Amide I region alone (**Supplementary S11**). This suggests that both wavelengths induce damage by similar mechanisms. Key peaks in the loadings spectra that were associated with the native protein include: 933 and 1296cm^-1^ (α-helix), 995 and 1243cm^-1^ (β-sheet), and 758, 869, 1010, 1216, 1576 and 1615cm^-1^ (aromatics). Key peaks in the loadings spectra that were associated with UVC irradiation included 983, 1268, 1665 and 1685cm^-1^ (nonregular structure) (*47*). We next included all spectra for both 227nm and 266nm irradiated rSARS-CoV-2 S in the same PCA. Across PC1 (dose-dependence of UVC radiation) 227nm irradiated rSARS-CoV-2 S spectra have larger scores than 266nm (**Supplementary S11**). From the above analysis we conclude that while both wavelengths have a similar mechanism of action, the effect of 227nm radiation is stronger, particularly on secondary structure. It is possible that the damage to aromatic residues and disulphide linkage, which are similarly effected by both 227nm and 266nm wavelengths are the main contributors to loss of S proteins binding ability and may explain the large loss in binding capacity observed at 100mJ (**Figure 3**).

### UVC irradiation does not cause sSARS-CoV-2 S glycan loss

The S protein is known to be decorated with 22 N-glycans per subunit (*23, 48*), with glycosylation contributing to the stabilisation of the RBD conformation (*49*), We observed changes in vibrations that relate to the glycosylation status of rSARS-CoV-2 S, but many overlap with those originating from protein bonds (*50*). However, we tentatively assigned the dose-dependent decrease in the vibration ∼1465cm^-1^ to glycan CH_2_ deformation and their degradation (**Supplementary S12**). SDS-PAGE analysis showed that UVC irradiation did not reduce glycosylation, but instead reduced Coomassie binding due to protein degradation.

## Conclusions

Here we have demonstrated the inactivation of SARS-CoV-2 by 266nm UVC, which matches closely with the absorption spectra of RNA and aromatic amino acids. 266nm light caused RNA damage at low powers, and we show in detail the mechanisms by which 266nm irradiation also damages conformation in recombinant SARS-CoV-2 spike protein through the cleavage of disulphide bonds and degradation of aromatic amino acids, reducing its ability to bind hACE2. Importantly we also investigate the effectiveness of 227nm which is well matched to protein backbone absorption. It was more effective at generating protein damage through oxidation and the unfolding of secondary structure. Reduced rSARS-CoV-2 S binding of hACE2 is observed as a result of UVC exposure and correlates well with changes in hydrophobicity and conformation. As expected, 227nm radiation was less effective at inducing RNA damage. Notably SARS-CoV-2 has among the largest of genomes for RNA viruses (*51*), making it especially sensitive to genomic damage. Therefore, the role of protein damage may be of even greater importance against other pathogens with smaller genomes. We therefore suggest that both wavelengths, leveraging dual inactivation mechanisms, could be more effective in preventing infectivity of viruses.

The results of inactivation studies with SARS-CoV-2 virus indicate a dose dependent inactivation but at doses that are >11.7 times higher than anticipated by our RNA damage assay, and 5-10 times lower than required to prevent recombinant spike protein binding hACE2. In addressing this difference we note that the ability of host cell ribosomes to process damaged RNA may differ from that of reverse transcriptase used in our assay. In addition, the recombinant spike protein used in the binding assays is more stabilised likely due to additional interactions compared to the wild-type spike protein. Further, the binding assay used does not detect the ability of the S protein to facilitate cell fusion, which likely also declines along with loss of hACE2 binding, or of damage to other viral proteins that may contribute to inactivation. Therefore, the overall effect of UVC protein damage on viral inactivation is likely to be underestimated.

Nonetheless we suggest a hierarchy of sensitivity to UVC irradiation with genomic damage preceding viral protein deactivation. We also note that for viruses with smaller genomes the relative contribution of RNA damage will decline increasing the importance of proteins as targets for UVC inactivation.

In summary we reveal dosages and mechanisms of SARS-CoV-2 inactivation that will also be applicable to other pathogens. Our work provides fundamental evidence that helps understand molecular targets of UVC wavelengths and dosage requirements for high throughput disinfection systems and devices to prevent the transmission and spread of airborne diseases, including Covid-19.

## ACKNOWLEDGEMENTS

We thank Jenny Russell (CL3 Lab Manager) and Mark Dixon (Life Sciences B85 Lab Manager) for facilitating the experiments and James Read and Aikta Sharma for help with analysis. We thank M Squared Lasers for providing the lasers on loan.

## AUTHOR CONTRIBUTIONS

GM, NHempler, RF and SM conceptualised the overarching collaborative project on laser disinfection. SM was the chief investigator, led the overall research team, coordinated and managed the research study. SM and SL led the design of experiments and analysis together with BS, ND, MC and CJM. MC, ND and CJM were principal investigators of the Spike protein, virus methodology development and SARS-CoV-2 virus work, respectively. NHanrahan and SM performed UVC irradiation experiments, UVC laser expertise was contributed by NH, RF and GM. GD, SM and PBJ performed UV-vis absorption spectroscopy experiments. GD performed Raman spectroscopy, Bis-ANS fluorescence and AFM experiments and analyzed data. BS, MCV and SL performed RNA, virus methodology development and PCR experiments and analysed data. JA expressed and purified proteins. JA performed SPR experiments and analyzed data. MVH, CS, RH, KS and CJM performed the SARS-CoV-2 work in CL3 labs, NHanrahan, SL and SM controlled the laser. JJW performed genome-scaling simulations for predicting required inactivation doses. G.D, P.B.J, NHanrahan and SL wrote the first draft of the manuscript with inputs from all authors. SL, GD, MC, CJM and SM led the finalisation of the manuscript. All authors contributed and approved the final version.

## Supplementary Information

### I. Materials and Methods

#### UV lasers and illumination through liquid light guide

The light source used for the experiments was based on a high-brightness, single-frequency, continuous-wave, solid-state laser operating at 532nm with a maximum output power of 20W (Equinox, M2 Lasers, Glasgow, UK). The 532nm light was doubled in frequency to 266nm by using a commercial cavity-enhanced second harmonic generation module (ECD-X-Q, M2 Lasers, Glasgow, UK). For the purpose of the experiments, the output power of the 532nm laser was limited to 2W, resulting in up to 468mW of 266nm light.

For the 227nm light, the same 532nm solid-state laser was used, however, the output was first used to pump a commercially available, highly frequency agile titanium sapphire laser (SolsTiS, M2 Lasers, Glasgow) that provided up to 6W of output power.

The laser and nonlinear conversion module were mounted on an optical breadboard for mobility, which was placed on either an optical bench during initial laboratory testing, or on a compact mobile trolley for working in a Containment Level 3 laboratory. The UV laser output passed through an optical shutter (SH1/M, Thorlabs) and was coupled into a 5mm core liquid light guide (LLG) (LLG5-4T, Thorlabs, UK) using two aluminium mirrors (PF10-03-G01, Thorlabs). The flexible LLG delivered high-power UV to the sample surface, and UV emission from the LLG was a uniform circle which diverged slowly. At 20mm from the sample surface, the UV illumination area covered a 23mm diameter circle. Exposure times were controlled using the electronic shutter (**Supplementary Figure S13**).

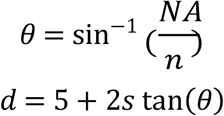

d = Spot size; s = Vertical distance to sample surface; NA = 0.42; A = LLG aperture size (5 mm)

#### UV illumination of liquid samples

A liquid sample (1-15μL) was pipetted onto the smooth upper surface of the central PMMA pedestal, and a quartz slide (UQG optics) was placed on top, providing a uniform thickness across the sample for UV exposure. The sample was illuminated with UV from above, through the quartz slide. After exposure, samples were recovered from the quartz and PMMA surfaces. More than 90% of the sample was typically recovered. UV transmission through the quartz slide was >92% at 266nm, and >86% at 227nm. Quartz slides were stored in 70% ethanol when not in use, and the quartz slides and sample chambers were cleaned with 70% ethanol between each sample irradiation.

#### UV transmission through liquids

Culture media (2mL) was placed in a quartz cuvette, aligned perpendicular to the beam propagation direction, and UV power transmission was measured through the filled cuvette with a power meter (30A-BB-18, Newport). 1mL of media was removed and replaced with 1mL DI water, and UV power transmission was measured again. This was repeated for sequential 2x dilutions of media with DI water to a 2^-15^ concentration. This was performed for DMEM supplemented with 0%, 4% or 10% FBS (**Supplementary Figure S14)** yielding molar extinction coefficients (mol^-1^.cm^3^).cm^-1^ for 266nm: 10%, 7.9; 4%, 4.8; 0%, 2.8 and 227nm: 10%, 48.3; 4%, 15.8; 0%, 7.4.

#### SARS-CoV-2 Virus and associated cell lines

Human coronavirus SARS-CoV-2 (BetaCoV/Australia/VIC01/2020) and a kidney cell line (Vero E6) were obtained from Public Health England (PHE), UK. Uninfected cells were maintained in DMEM (Invitrogen; cat no 11995065) supplemented with 10% foetal bovine serum (FBS), glutamine and 50u/ml penicillin-streptomycin at 37°C in 5% CO_2_. To produce working stocks of virus, cells were infected at a multiplicity of infection of 0.005 then maintained for 3 days in DMEM supplemented with 4% foetal bovine serum, 50U/mL penicillin-streptomycin and 25mM HEPES. The resultant cell culture supernatant was clarified by centrifugation at 2,000g for 10 minutes and frozen as aliquots at -80°C.

All handling of live and inactivated SARS-CoV-2 virus was performed in containment level 3 laboratories, with dried samples being handled in a class II microbiological safety cabinet (MSC) and liquid samples in a class III MSC.

#### UV inactivation assay for SARS -CoV-2

25μL microliters of SARS-CoV-2 (1×10^6^ pfu.mL^-1^) were placed in the centre of alternate wells of a 24-well plate and left to dry under ambient conditions for a period of 3 hours. Laser power from the LLG was set for continuous exposure at specified output power. Dried virus samples were illuminated with UV at 266nm for 1, 3, 10, 30, and 100s, and all exposures were repeated in triplicate. Each exposed sample was resuspended in 0.5ml serum-free DMEM supplemented with 25mM HEPES (infection medium) for a period of 30 minutes and the samples and their respective dilutions assessed by plaque assay.

#### SARS-CoV-2 plaque assay

Vero E6 cells were seeded at 2.5×10^5^ cells/well in a 12-well plate and left for a period of 24 hours. Cells were washed once with infection medium and 0.4ml virus-containing supernatants added to each well. After a 1 hour incubation at 37°C in 5% CO_2_, infectious supernatants were removed and a 1.5mL overlay of 1 x DMEM supplemented with 4% FBS, 25mM HEPES and 0.6% (w/v) cellulose (Sigma; cat no 435244) was added. Plates were incubated at 37°C and 5% CO_2_ for 72 hours before removing the overlay, fixing with 8% formaldehyde in PBS, and staining with 0.1%(w/v) crystal violet in a solution of 20% (v/v) ethanol.

#### RNA damage assay

MS2 phage ssRNA (Sigmal Aldrich) was resuspended at a concentration of 80ng/uL in nuclease-free water and 15μL drops placed in the lid of a sterile 96-well plate were irradiated with 21mW power using 227nm or 266nm lasers, for 0, 1, 3, 10, 30 or 100s. 10 μL of RNA solution was recovered (800ng RNA) and reverse transcribed using a Precision Nanoscript2 RT kit (Primer Design Ltd) using a first strand synthesis primer (CCAAATCGGGAGAATCCCGGGTCC). 10ng of the equivalent cDNA generated was then subjected to qPCR using a qPCRBio SyGreen Kit (PCR Biosystems Ltd) using two primer pairs situated 767-925 bases (ATCCGCTCGCACTACGGAAT, ATGCCTATGGTTCCGGCGTT) or 2087-2231 bases (AGAGCCCTCAACCGGAGTTT, TAAGCCTGTGAGCGCGAGTT) from the 5’ end of the first strand cDNA. 40 cycles of denaturation at 95°C, 5s; and annealing/extension at 62°C, 30s; were performed using StepOne Plus machine (Applied BioSystems). Each reaction was carried out in duplicate and deltaCt values were calculated and normalised against non-irradiated controls. Curve fitting analysis was performed using prism8 (Graphpad Software LLC). 99.9% inactivation doses were calculated from the fitted curve.

#### UV dose-dependent lesion scaling with ssRNA genome length

Considering light interacting with RNA genomes, replication inhibition by UVC lesion effects were modelled using Poisson statistics involving a linear proportionality relating to the genome length. For predicting wavelength dependent doses required to inactivate 99.9% of a virus with a given ssRNA genome (e.g. Influenza A, 13,588 nt; and SARS-CoV-2, 30,000nt). The probability model was normalized using the experimental replication inhibition values obtained with 21mJ/cm^2^ UV doses interacting with the MS2 2,231 base fragment.

#### Expression and purification of trimeric recombinant SARS-CoV-2 spike protein

To express the prefusion S ectodomain, a gene encoding residues 1−1208 of SARS-CoV-2 S (GenBank: MN908947) with proline substitutions at residues 986 and 987, a “GSAS” substitution at the furin cleavage site (residues 682–685), a C-terminal T4 fibritin trimerization motif, an HRV3C protease cleavage site, a TwinStrepTag and an 8XHisTag was synthesized and cloned into the mammalian expression vector pαH. Expression plasmid encoding SARS-CoV-2 S glycoprotein 6 was transiently transfected into Human Embryonic Kidney (HEK) 293F cells. Cells were maintained at a density of 0.2-3×10^6^ cells per mL at 37°C, 8% CO_2_ and 125rpm shaking in FreeStyle 293F media (Fisher Scientific). Prior to transfection two solutions containing 25mL Opti-MEM (Fisher Scientific) medium were prepared. Plasmid DNA was added to one to give a final concentration after transfection of 310μg/L. Polyethylenimine (PEI) max reagent (1mg/mL, pH 7) was added to the second solution to give a ratio of 3:1 PEI max: plasmid DNA. The two solutions were combined and incubated for 30 minutes at room temperature. Cells were transfected at a density of 1×10^6^ cells per ml and incubated for 7 days at 37°C with 8% CO_2_ and 125 rpm shaking.

After harvesting, the cells were spun down at 4000rpm for 30 minutes and the supernatant applied to a 500mL Stericup-HV sterile vacuum filtration system (Merck) with a pore size of 0.22μm. The supernatant containing SARS-CoV-2 S protein was purified using 5mL HisTrap FF column connected to an Akta Pure system (GE Healthcare). Prior to loading the sample, the column was washed with 10 column volumes of washing buffer (50mM Na_2_PO_4_, 300mM NaCl) at pH 7. The sample was loaded onto the column at a speed of 2mL/min. The column was washed with washing buffer (10 column volumes) containing 50mM imidazole and eluted in 3 column volumes of elution buffer (300mM imidazole in washing buffer). The elution was concentrated by a Vivaspin column (100kDa cut-off) to a volume of 1mL and buffer exchanged to phosphate buffered saline (PBS).

The Superdex 200 16 600 column was washed with PBS at a rate of 1mL/min. After 2 hours, 1mL of the nickel affinity purified material was injected into the column. Fractions separated by SEC were pooled according to their corresponding peaks on the Size Exclusion chromatograms. The target fraction was concentrated in 100kDa vivaspin (GE healthcare) tubes to ∼1mL.

#### Expression and purification of hACE2

FreeStyle293F cells (Thermo Fisher) were transfected with polyetyhlenimine and a plasmid encoding residues 1-615 of human ACE2 with a C-terminal HRV3C protease cleavage site, a TwinStrep Tag and an 8XHisTag. This construct is identical to full length ACE2 except is truncated at position 626. This protein was expressed and purified identically as for the hSARS-CoV2 S glycoprotein, with the exception of a smaller Vivaspin cutoff being used for buffer exchanging.

#### His-Tag removal of hACE2

Following purification, the His-Tag was removed from hACE2 using HRV3C protease cleavage (Thermo Fisher). Digestion was performed at a ratio of 1:20 HRV3C protease: ACE2 in 1x HRV3C reaction buffer (Thermo Fisher) and incubated at 4°C overnight. To remove the HRV3C and uncleaved hACE2 nickel affinity chromatography was performed, except the flow through was collected rather than the elution.

#### Absorbance Spectroscopy

Absorbance spectroscopy was performed using a NanoDrop 2000 spectrometer. Spectra were collected between 220nm and 340nm using a 2μl sample, all spectra were collected immediately after UV irradiation to minimise contributions from aggregation. Data was extracted from an average of 3 repeat readings, no further processing was performed. Erroneous data was rejected if spectra were flat or unusually noisy, typically due to the presence of an air bubble during data collection. Nonlinear curve fit analysis was performed using Origin 2020.

#### Tryptophan fluorescence

Tryptophan fluorescence was performed using a Cary Eclipse fluorescence spectrophotometer, excitation was set to 280nm and an emission collected between 300nm and 500nm, excitation and emission slit widths were set to 5nm. Protein concentration was 9μg/mL in all cases. Displayed spectra are blank subtracted averages of 3 repeat readings and have undergone nearest neighbour averaging with a 5-point window.

#### Bis-ANS fluorescence

Bis-ANS (Sigma) was dissolved in PBS and mixed with protein samples after irradiation to a final concentration of 10μM dye and 0.4mg/ml protein. Solutions were transferred to 96-well plates (Costar) and fluorescence was measured using an excitation filter of 450nm and an emission filter of 490nm. Each experiment was performed using two technical duplicates. The fluorescence of each equivalent blank solution (buffer + dye) was subtracted from each experimental readout and fluorescence was normalized as fold change over native rSARS-CoV-2 S protein fluorescence.

#### Atomic force microscopy (AFM)

20μL of *2*ug/mL protein was added to a freshly cleaved 10mm mica disc (Agar Scientific) and incubated at room temperature for 2 min. The protein solution was then washed with 0.22μM filtered, double distilled H_2_O three times before drying in air. Samples were imaged using a Digital Instruments Multimode IV AFM system operated in tapping mode. Aluminum-coated, noncontact/tapping mode probes with a resonance frequency of 320kHz and force constant of 42N/m were used for all images (Nanoworld, POINTPROBE NHCR). Probes were autotuned using Nanoscope III 5.12r3 software before use. Images shown are representative of the sample. Two 5μm images were taken at random points on the sample per experiment with a scan rate of 1–2Hz and 512 samples per line/512 lines per image. Height analysis was performed using the particle analysis feature in WSxM Beta (*52*) using 10 bins with a range of 0-50nm.

#### Raman spectroscopy

A Renishaw InVia microscope system was used for Raman spectroscopy. Quartz coverslips were coated in trichloro(1H,1H,2H,2H-perfluoroocytl)silane (Sigma) by chemical vapor deposition. A silane atmosphere was created by a 30 min desiccation using a small volume of trichloro(1H,1H,2H,2H-perfluoroocytl)silane in a reaction chamber. Quartz coverslip surfaces were activated by oxygen plasma treatment before incubation in the silane atmosphere to create a silane monolayer on the coverslip surface for 2h. For Drop-deposition Raman spectroscopy (DDRS), 0.5μL of each protein sample was first dried onto a quartz coverslip for 12 min under a vacuum. Spectra were then collected from the “coffee ring” of each drop, where proteins are found in the absence of bulk salt (*47, 53, 54*). The samples were excited using a 785nm laser focused through a Leica 50× (0.75 NA) short working distance objective for DDRS. Data was obtained and parameters were set using Renishaw WIRE4.1 software, WIRE was also used for cosmic ray removal. Spectra were collected in the fingerprint region (614–1722cm^−1^). The Raman system was calibrated to the 520cm^−1^ reference peak of silicon prior to each experiment.

Spectra were acquired for 5s with 6 acquisitions in different locations on the coffee ring per experiment. A total of 15 spectra were collected for native rSARS-CoV-2 S and 11 spectra were collected for each UVC condition. Erroneous spectra were rejected with unusual background fluorescence that could not be removed using polynomial subtraction. Presented Spectra represent the class means of all remaining spectra. Preprocessing and Principal Component Analysis (PCA) was performed using the IRootLab plugin (0.15.07.09-v) for MATLAB R2015a (*55*). All spectra were background-subtracted using blank quartz spectra and were smoothed using the wavelet denoising function. A fifth-order polynomial was used to remove fluorescence and the ends of each spectra were anchored to the axis using the rubberband-like function. Spectral intensity normalization was applied using vector normalisation or using a specific band (each mentioned within captions). Trained-mean centering was then applied to the spectra before PCA with a maximum of ten principle components. Amide I second derivative analysis was performed on class mean spectra using WIRE4.1 software.

#### Surface plasmon resonance (SPR)

rSARS-COV-2 S and hACE2 proteins were buffer exchanged in HBS P+ buffer (Cytiva/GE Healthcare). All analysis was performed using a Biacore T200 (Cytiva/GE Healthcare). After removing metallic contaminants via a pulse of EDTA (350 mM) for 1 min at a flow rate of 30μL/min, the chip was loaded with Ni_2_+by injecting NiCl_2_ for 1 min at a flow rate of 10μL/min. SARS-CoV2 S protein (50,000ng/mL) was injected at 10μL/min for 240s. Control channels received neither trimer nor NiCl_2_. Control cycles were performed by flowing the analyte over Ni_2_+-loaded NTA in the absence of trimer; there were no indications of non-specific binding. The analyte was injected into the trimer sample and control channels at a flow rate of 50μL/min. Serial dilutions ranging from 200nM to 3.125nM were performed in triplicate along with HBS P+ buffer only as a control. Association was recorded for 300s and dissociation for 600s. After each cycle of interaction, the NTA-chip surface was regenerated with a pulse of EDTA (350mM) for 1 min at a flow rate of 30μL/min. A high flow rate of analyte solution (50μL/min) was used to minimize mass-transport limitation. The resulting data were fit to a 1:1 binding model using Biacore Evaluation Software (GE Healthcare) and these fitted curves were used to calculate KD.

#### Custom sample holders for UV illumination

Sample holders were created from PMMA, a microscope slide, and a quartz slide for illumination of a known liquid sample thickness. PMMA (3mm, Techsoft) was cut into 10mm squares using a laser cutter. Further pieces of PMMA were laser etched to remove a depth of 150μm, and then also cut to 10mm squares. Glass slides were also lightly etched to provide a high surface area for PMMA binding. The etched PMMA pedestal was attached to the centre of a microscope slide with epoxy resin, with the etched surface in contact with the slide. Two 3mm thick PMMA pedestals were similarly attached either side of the central pedestal, resulting in the central pedestal being 150μm lower than the surrounding supports. Final sample thickness was confirmed using digital callipers to be 150±5μm. Samples were placed on the central pedestal, and the quartz glass placed over, resting on the outer pedestals.

### II. Supplementary Table

**Supplementary Table 1:**
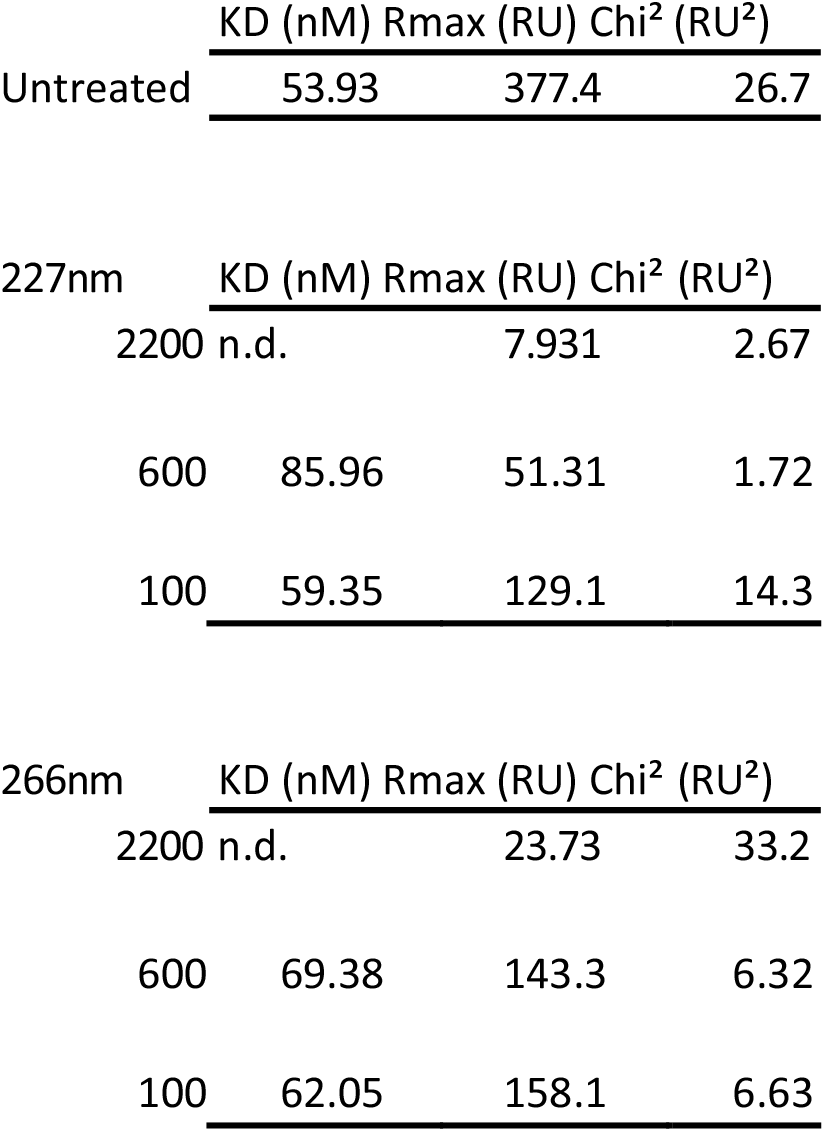
Parameters for SPR binding of SARS-CoV2 and hACE2.

### III. Supplementary Figures

**Supplementary Figure 1.**
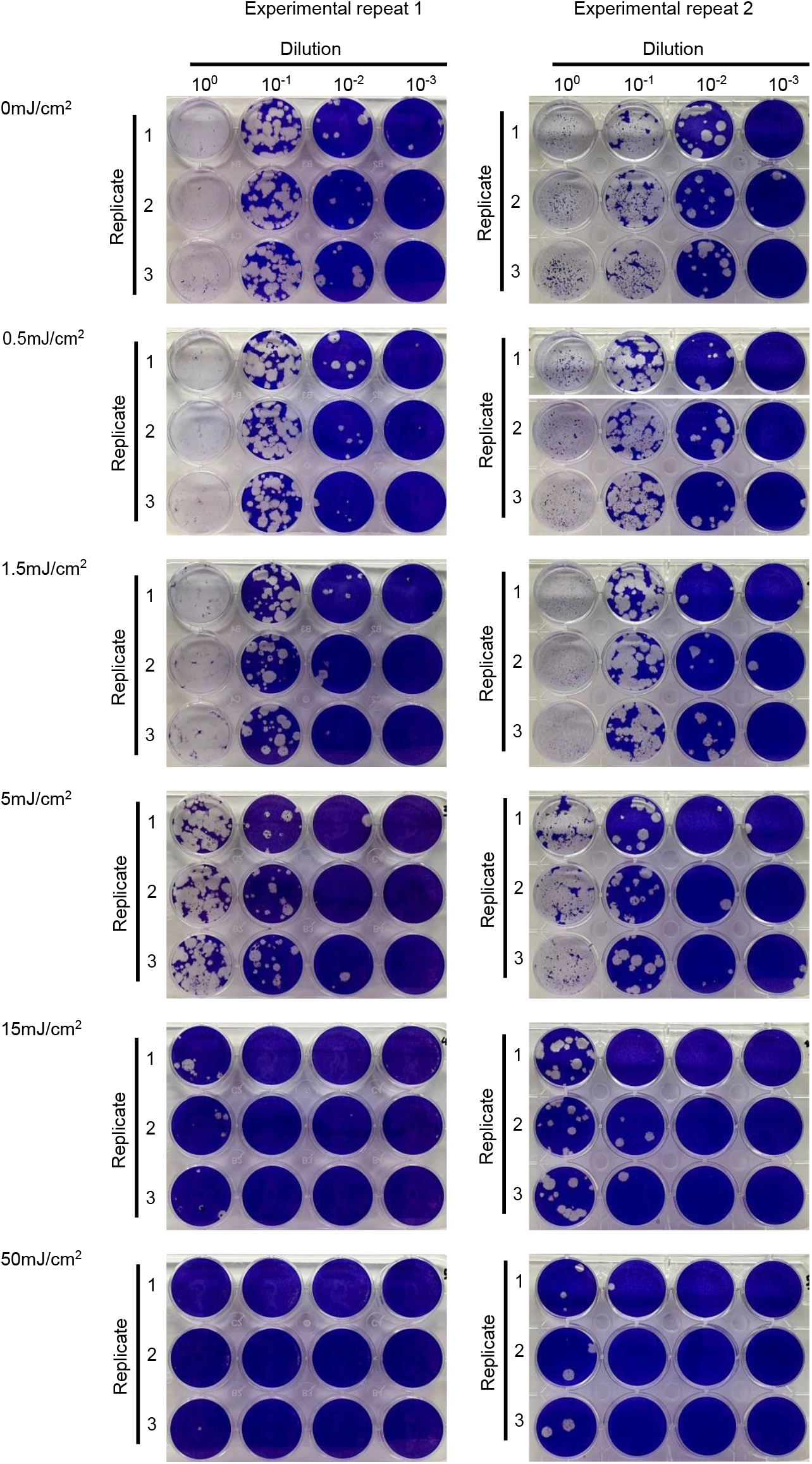
Plaque assay for SARS-CoV-2 virus inactivation after UV irradiation. SARS-CoV-2 virus was irradiated with 266nm light, with laser power density of 0.5mW/cm^2^ and exposure times of 1, 3, 10, 30 and 100s, repeated in triplicate. Cell monolayers were stained with crystal violet and fixed with PFA. Images show that with increasing UV dose, fewer plaques formed on cell monolayers infected with UV-irradiated virus stock. 99.9% of virus was inactivated with a dose of 50mJ/cm^2^.

**Supplementary Figure 2.**
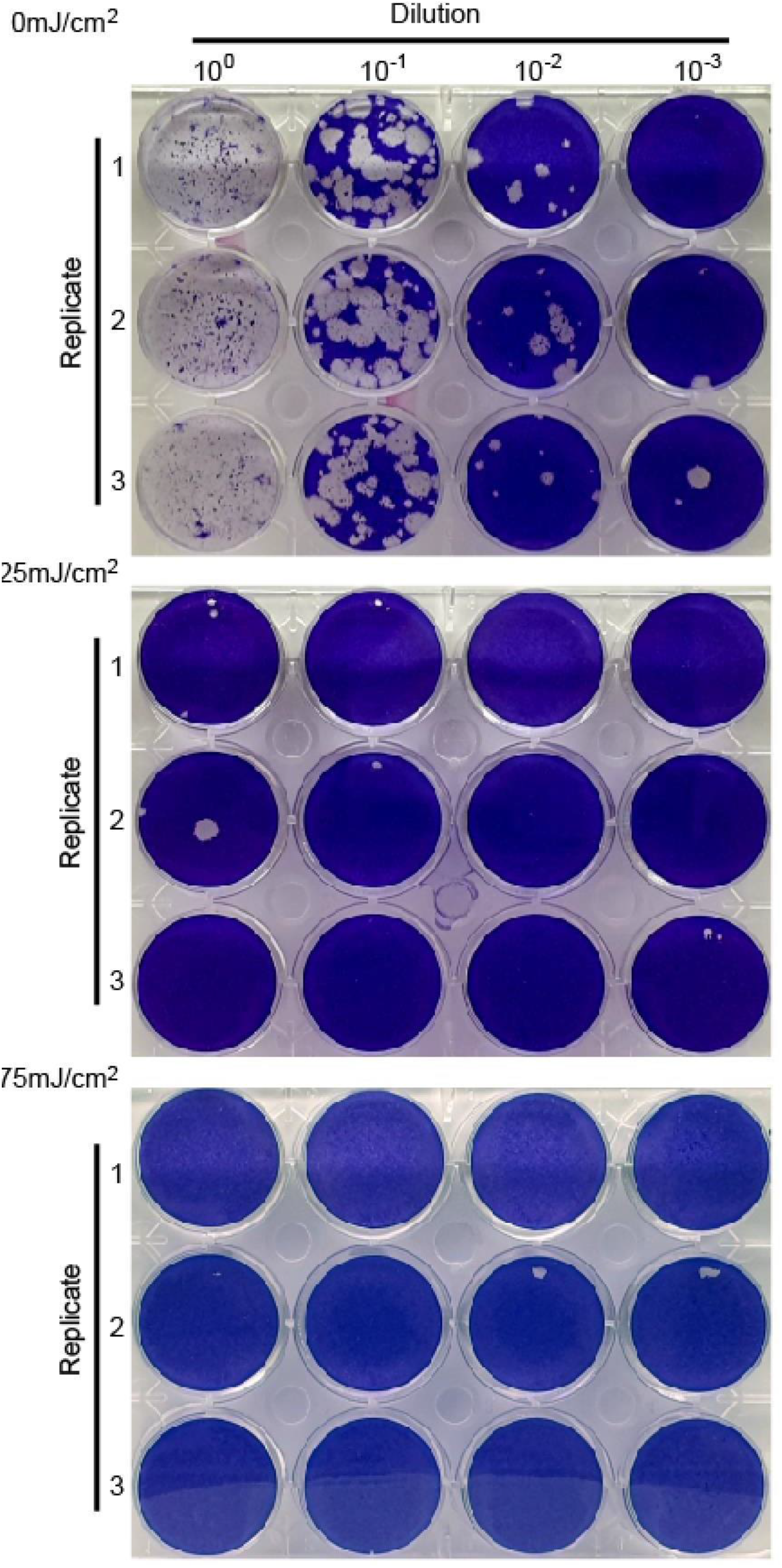
Plaque assay for SARS-C0V-2 virus inactivation after high dose UVC irradiation. SARS-CoV-2 virus was irradiated with UVC at 266nm, with laser power density of 25mW/cm^2^ and exposure times of 0, 1 3, 10, 30 and 100s, repeated in triplicate. Cell monolayers were stained with crystal violet and fixed with PFA. Images show a single plaque formed for only one of the three repeats at the lowest UVC dose (25mJ/cm^2^), and no plaques were observed for higher doses (10, 30 and 100s not shown), indicating complete inactivation of the virus.

**Supplementary Figure 3.**
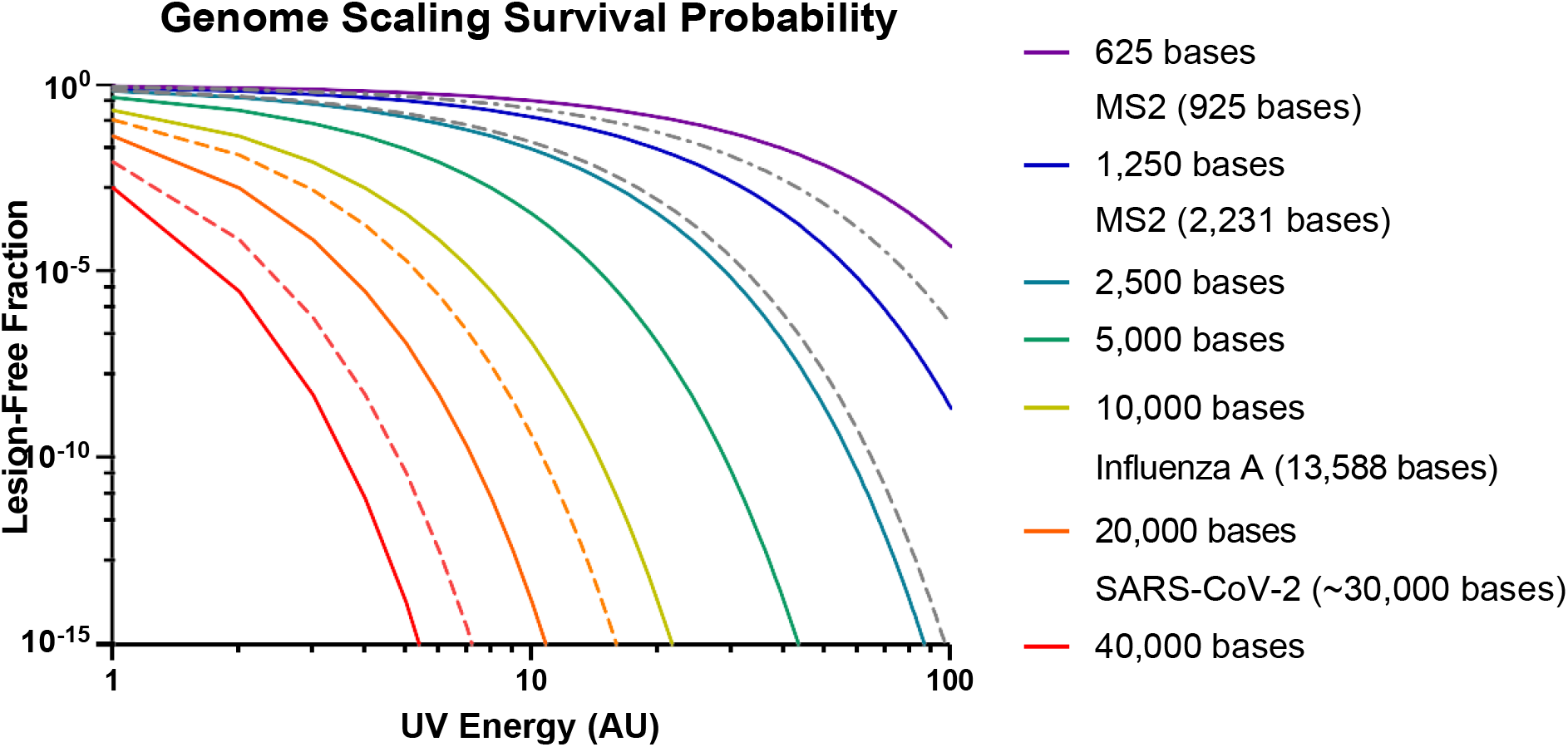
Theoretical genome scaling analysis. The proportion of RNA that remains lesion free plotted against Energy, for various sized genomes. The probability model was normalized using the experimental replication inhibition values obtained with 21mJ/cm^2^ UV doses interacting with the ∼2.3kb MS2 fragment.

**Supplementary Figure 4:**
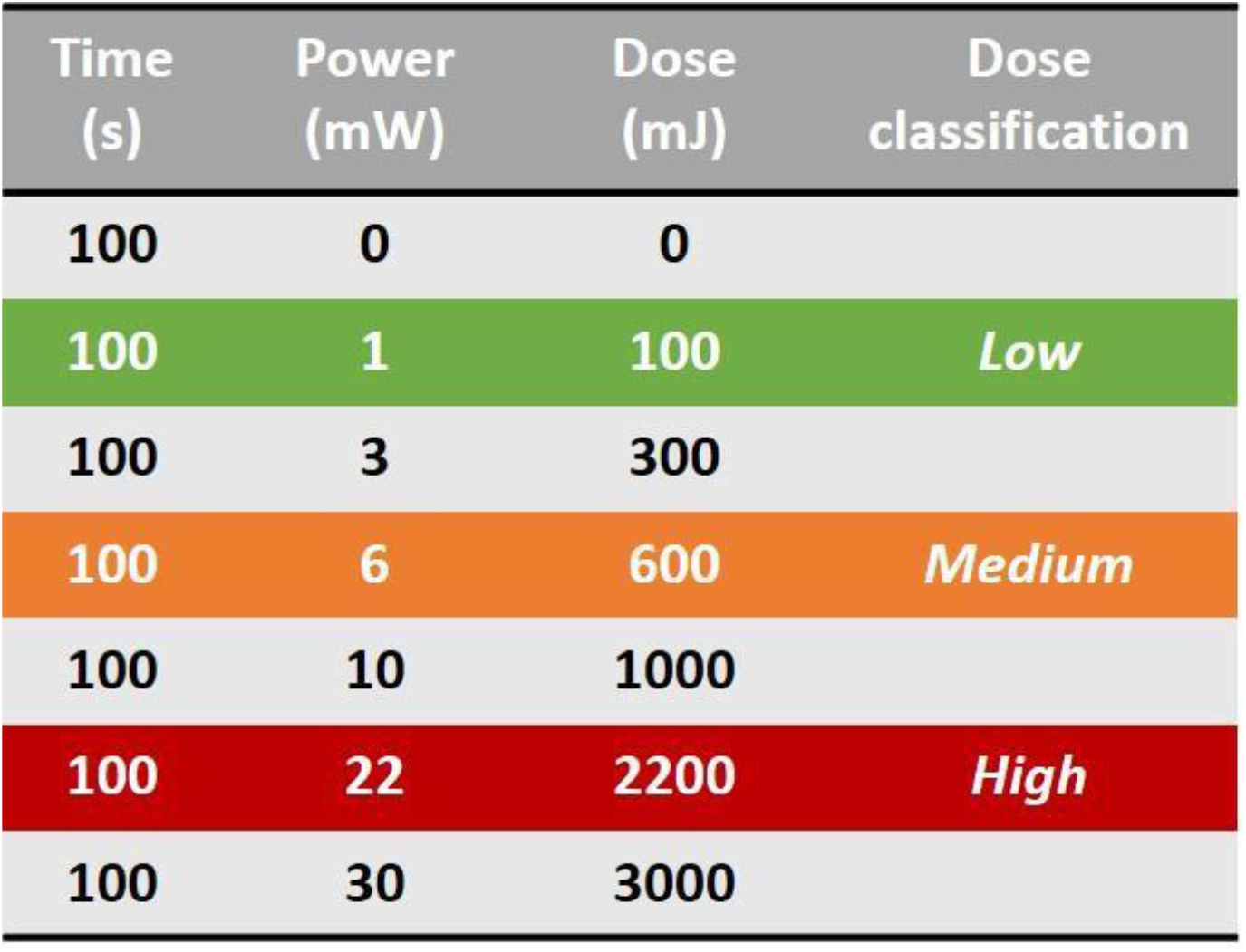
UVC doses used to irradiate rSARS-CoV-2 S protein. All doses shown were used for UV-vis absorption spectroscopy. Highlighted doses were used for experiments in figures 3, 4 and 5.

**Supplementary Figure 5.**
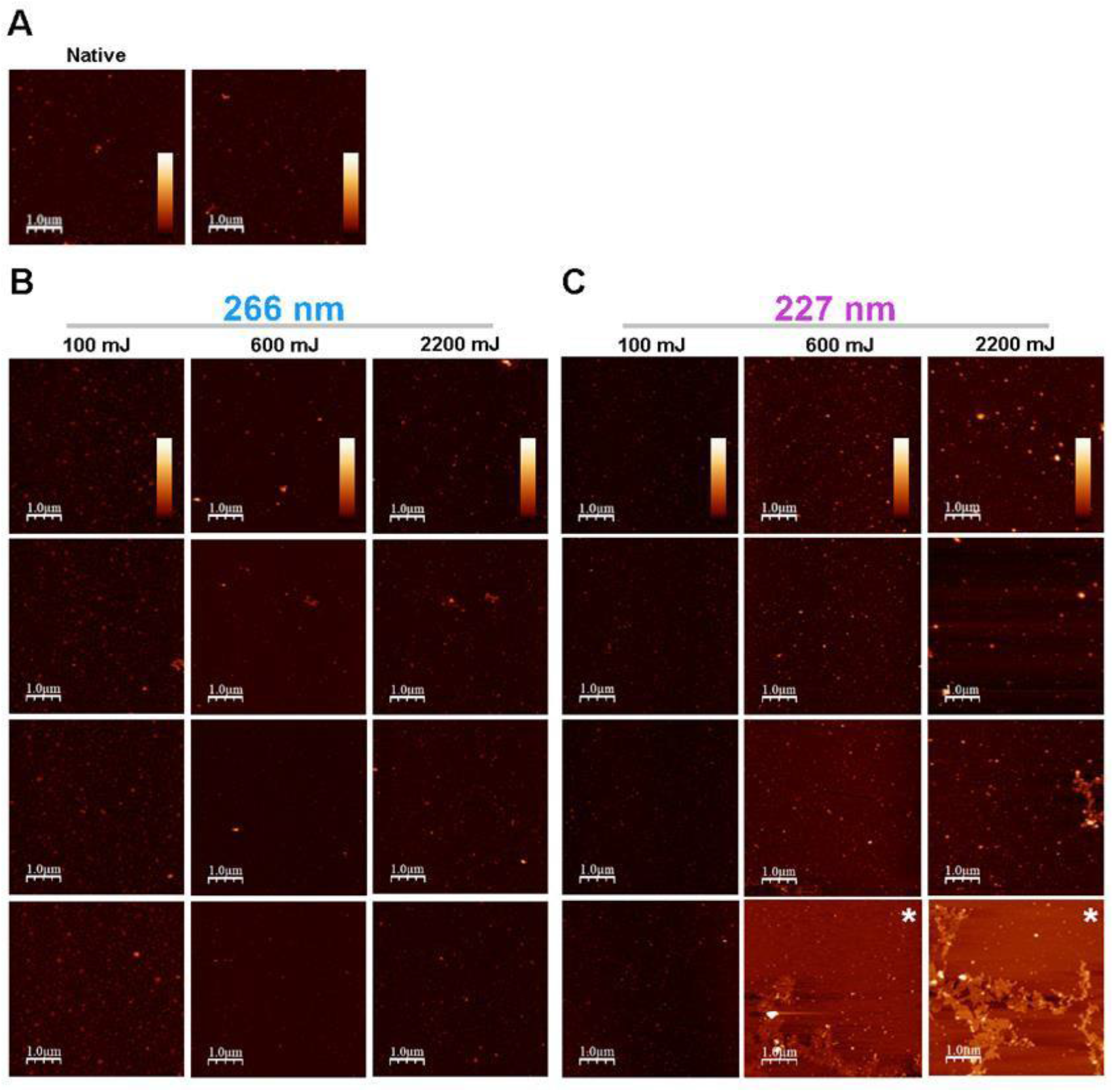
AFM images of SARS-CoV-2 S used for particle height analysis. AFM images used for particle height analysis shown in Figure 3. AFM images of SARS-CoV-2 S bound to a mica surface in the native from (A), or after UVC irradiation using 266nm light (B) or 227nm light (C). Images are 5μm x 5μm, scale bar is equal to 1μm and z-scale is equal to 0-30nm. Images labelled with * were not used for height analysis as large aggregated structures prevented correct image flattening resulting in incorrect Z-values.

**Supplementary Figure 6.**
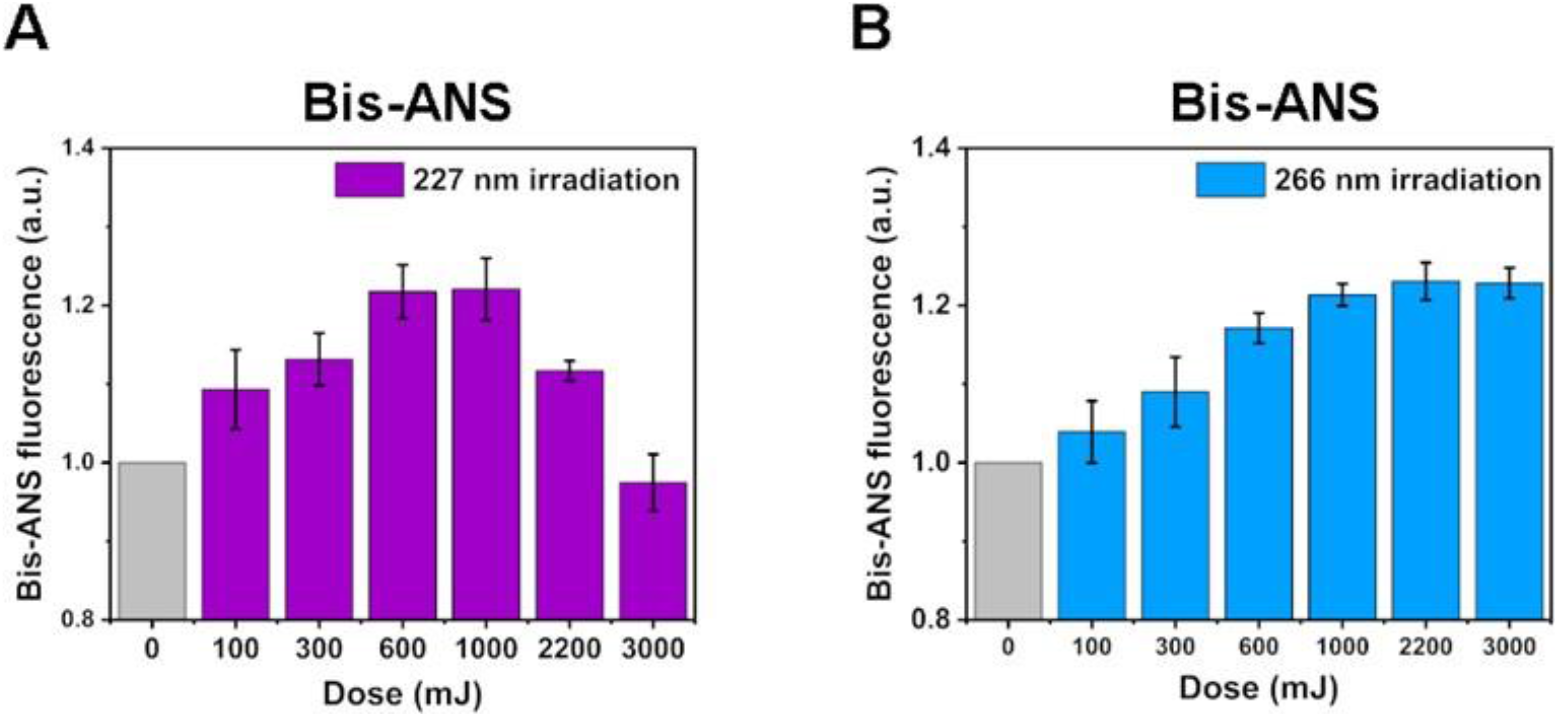
Dose-dependent changes in BSA surface hydrophobicity in response to UVC irradiation follow same pattern as for rSARS-CoV-2 S. Changes in Bis-ANS binding, measured by fluorescence emission at 490nm, induced by 227nm radiation (A) and 266nm radiation (B). n=2, the plotted data represent the average and SD from four fluorescence readings.

**Supplementary Figure 7.**
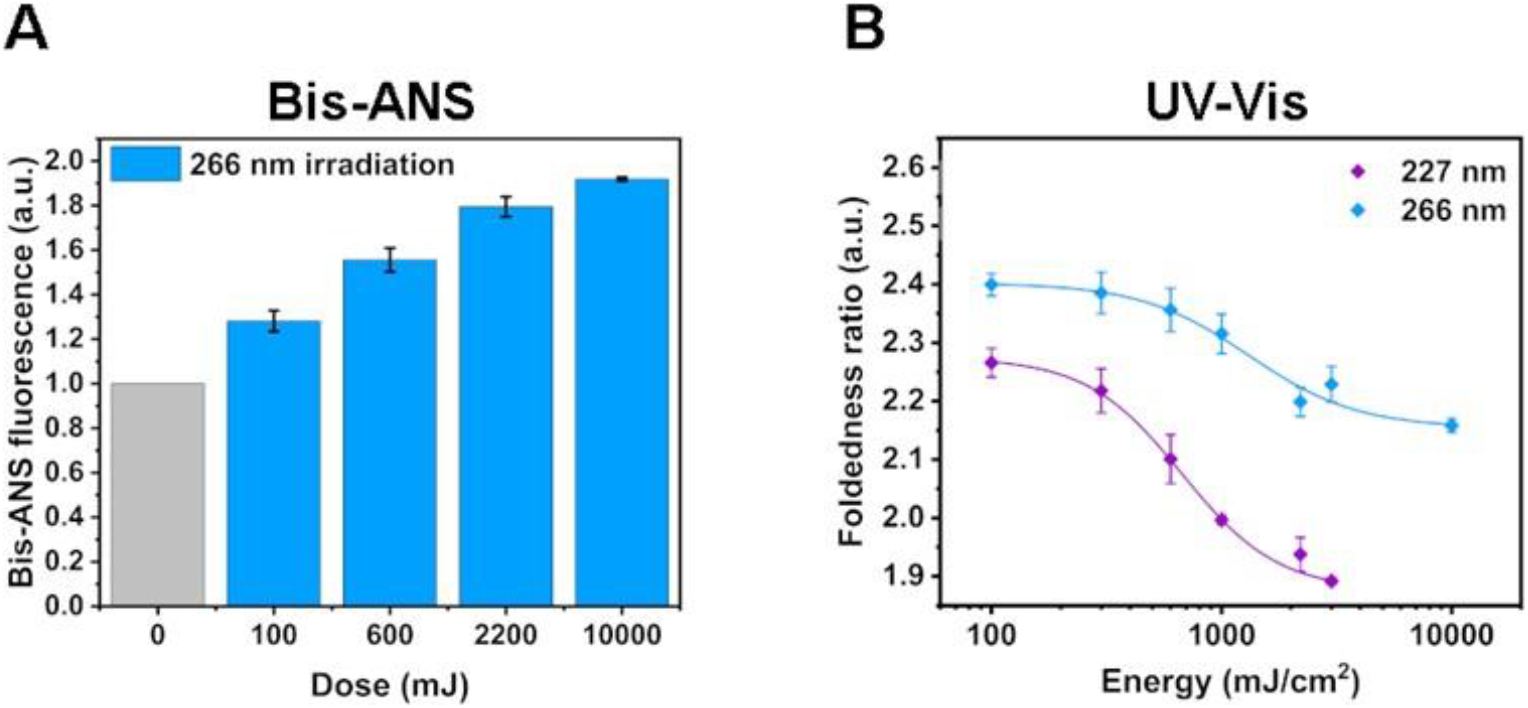
10J 266nm dose does not induce the same conformational effects in SARS-CoV-2 S as lower doses 227nm radiation. A) Changes in Bis-ANS binding measured by fluorescence emission at 490nm induced by 266nm radiation. B) Dose-dependance of rSARS-CoV-2 spike protein foldedness ratio to UVC radiation determined by UV-vis absorption spectroscopy. Foldedness ratio = A280nm/A275nm + A280nm/A258nm. n=1. The plotted data for 10000mJ represents the average and SD from 2 fluorescence/absorbance readings, all other plotted data represent the average and SD from 4 fluorescence/absorption readings (n=2).

**Supplementary Figure 8.**
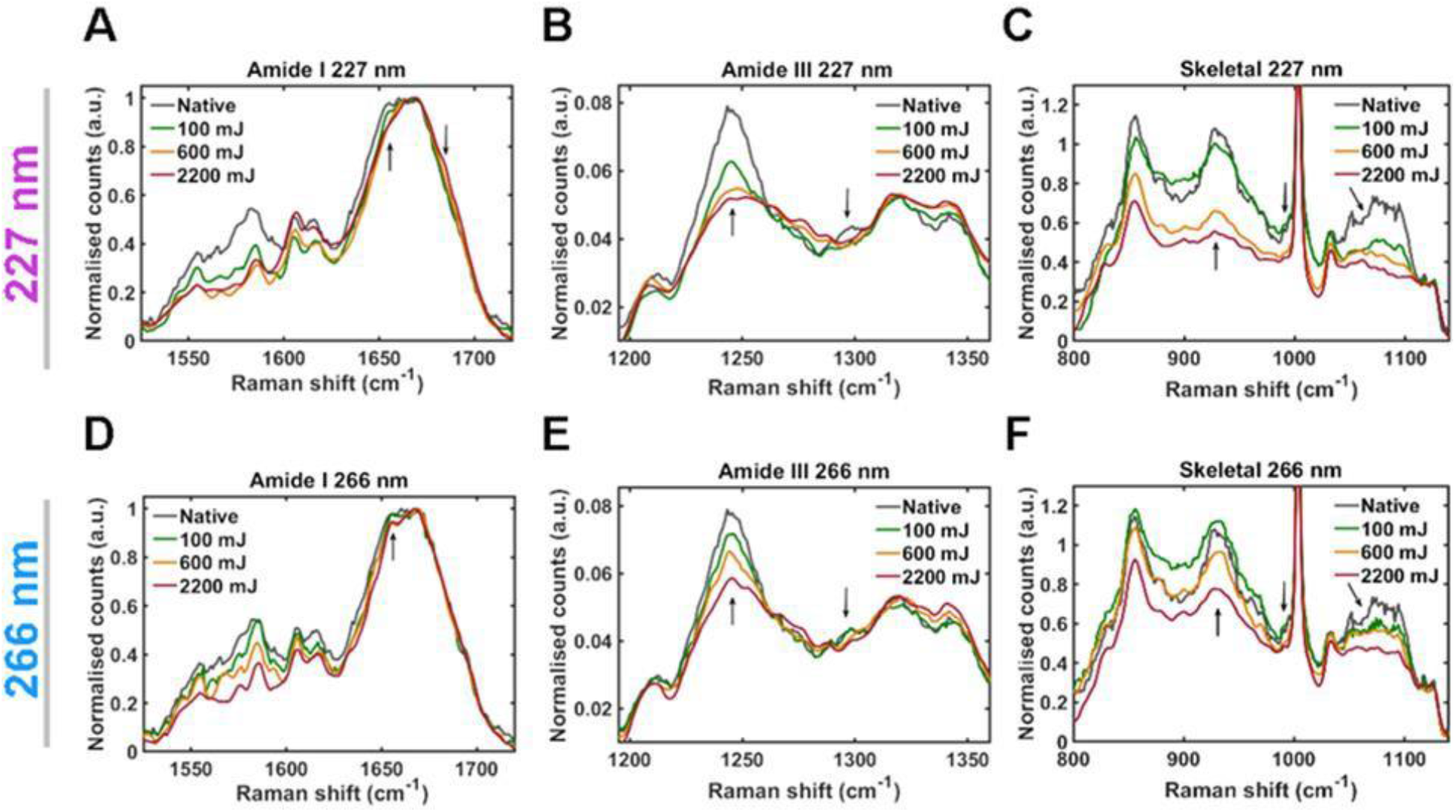
UVC dose-dependence of rSARS-CoV-2 spike protein secondary structure determined by Raman spectroscopy. Changes in Raman spectra induced by 227nm radiation (A-C) or 266nm (D-F). A, D) Amide I spectra from 1525-1722cm^-1^. Arrows indicate points of variation due to change of UVC dose. Upwards arrows indicate α-helix vibration, downwards arrow indicates nonregular structure vibration. B, E) Amide III spectra from 1195-1360cm^-1^. Upwards arrow indicates β-sheet and mannose vibrations, downwards arrow indicates α-helix vibration. C, F) Skeletal region spectra from 800-1140cm^-1^. Upwards arrow indicates α-helix vibration, downwards arrow indicates β-sheet and mannose vibration. n=2, plotted spectra represent the class means of 10-15 spectra per class.

**Supplementary Figure 9.**
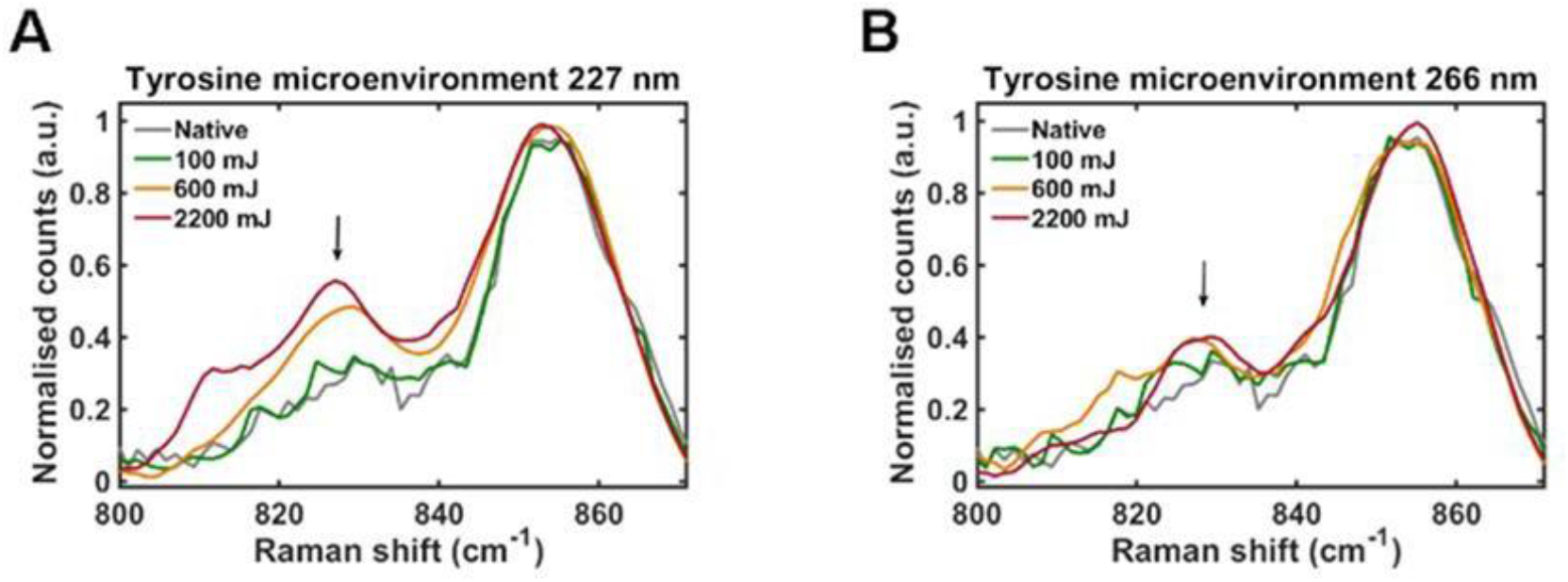
UVC dose-dependence of rSARS-CoV-2 spike protein tyrosine microenvironment determined by Raman spectroscopy. A) Changes in 830cm^-1^/850cm^-1^ ratio induced by 227nm radiation. B) Changes in 830cm^-1^/850cm^-1^ratio induced by 266nm radiation. Spectra are normalised to ∼850cm^-1^ tyrosine peak. Arrows indicate 830cm^-1^ tyrosine peak. n=2, plotted spectra represent the class means of 10-15 spectra per class.

**Supplementary Figure 10.**
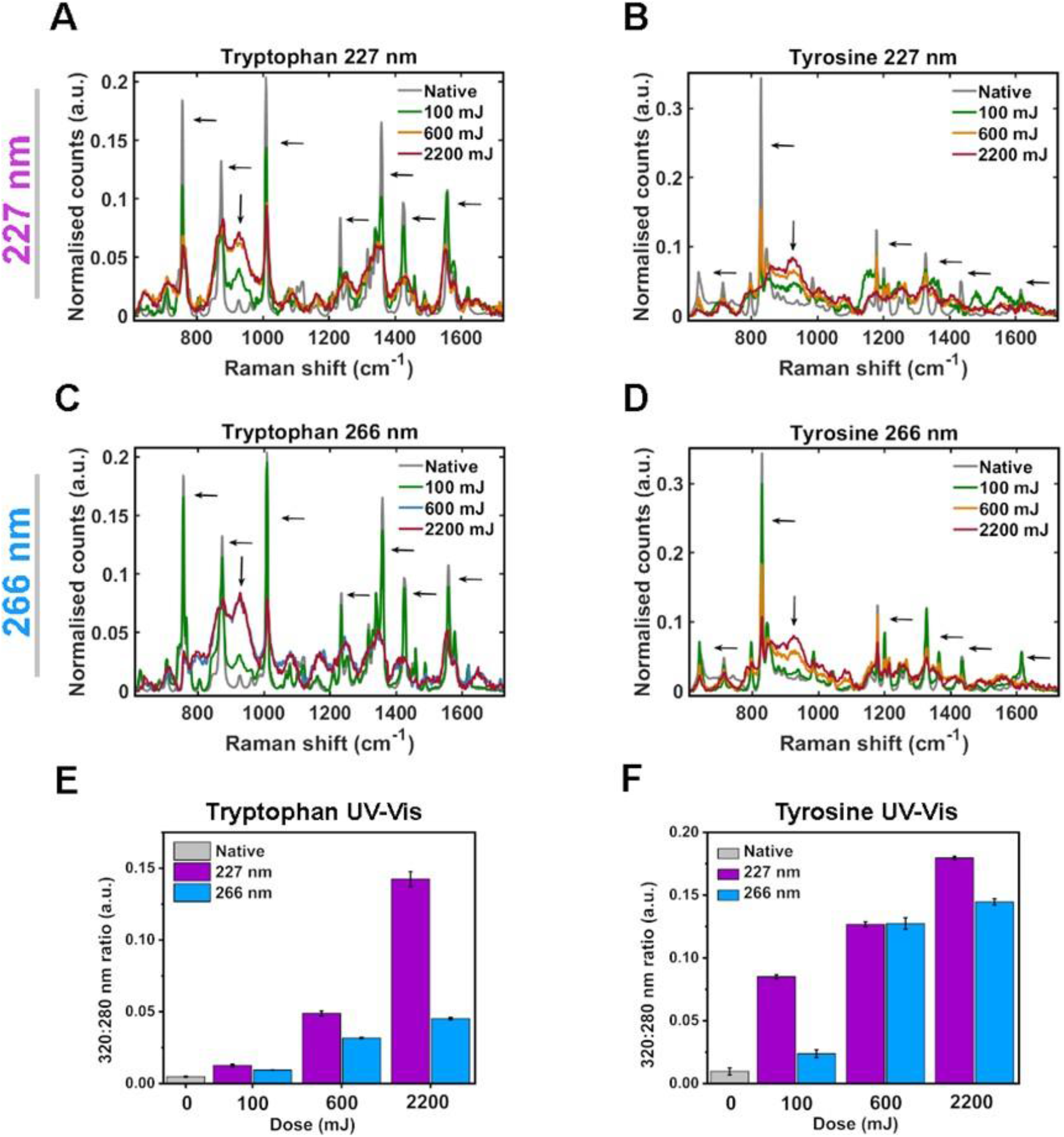
UVC dose-dependence degradation of pure aromatic amino acids. A-B) Changes in Raman fingerprint induced by 227nm radiation for tryptophan (A) and tyrosine (B). Downwards arrows indicate C-C vibrations and horizontal arrows indicate aromatic vibrations. C-D) Changes in Raman fingerprint induced by 266nm radiation for tryptophan (A) and tyrosine (B). Downwards arrows indicate C-C vibrations and horizontal arrows indicate aromatic vibrations. n=1, plotted spectra represent the class means of 3 spectra per class. E-F) Dose-dependance of tryptophan (E) and tyrosine (F) oxidation ratio to UVC radiation determined by UV-vis absorption spectroscopy. Oxidation ratio = A320/A280nm. n=1, the plotted data represent the average and SD from 3 absorbance readings per class.

**Supplementary Figure 11.**
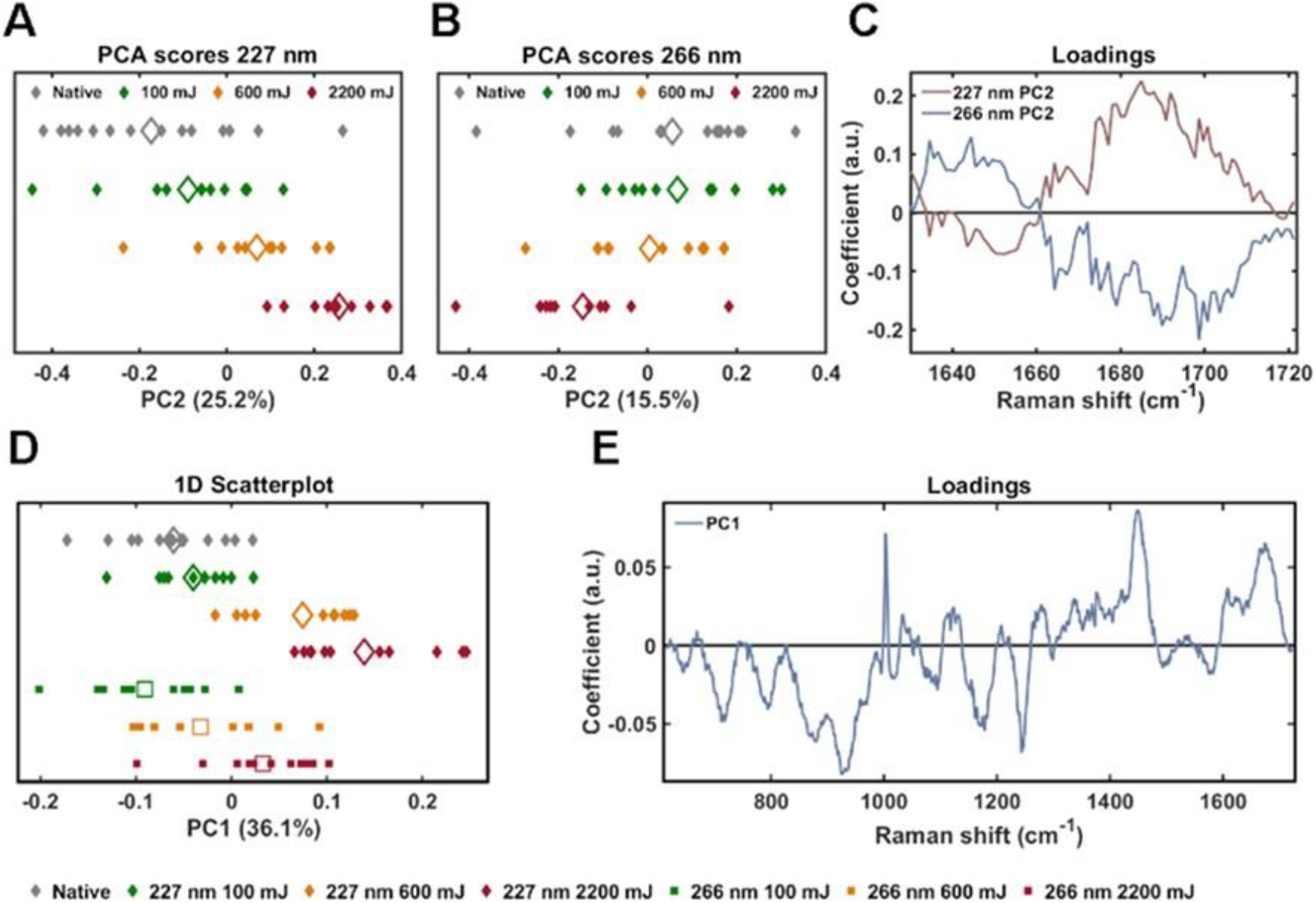
PCA of rSARS-CoV-2 S Raman spectra. A-B) 1-dimensional principle component analysis (PCA) scores plot of Amide I Raman spectra (1635-1722cm^-1^) for 227nm (A) and 266nm (B) irradiated rSARS-CoV-2 S. Each solid diamond represents the PC score of a single spectrum. Hollow diamonds represent mean score. C) PC loadings spectra representing the spectral variation responsible for the score across the given PC axis. D) 1-dimensional principle component analysis (PCA) scores plot of Raman spectra (622-1722cm^-1^) for 227nm and 266nm irradiated rSARS-CoV-2 S. Each solid diamond/square represents the PC score of a single spectrum. Hollow diamonds represent mean score. E) PC loadings spectra representing the spectral variation responsible for the score across the given PC axis. n=2, PCA performed on 10-15 spectra per class, 10 PCs were retained.

**Supplementary figure 12.**
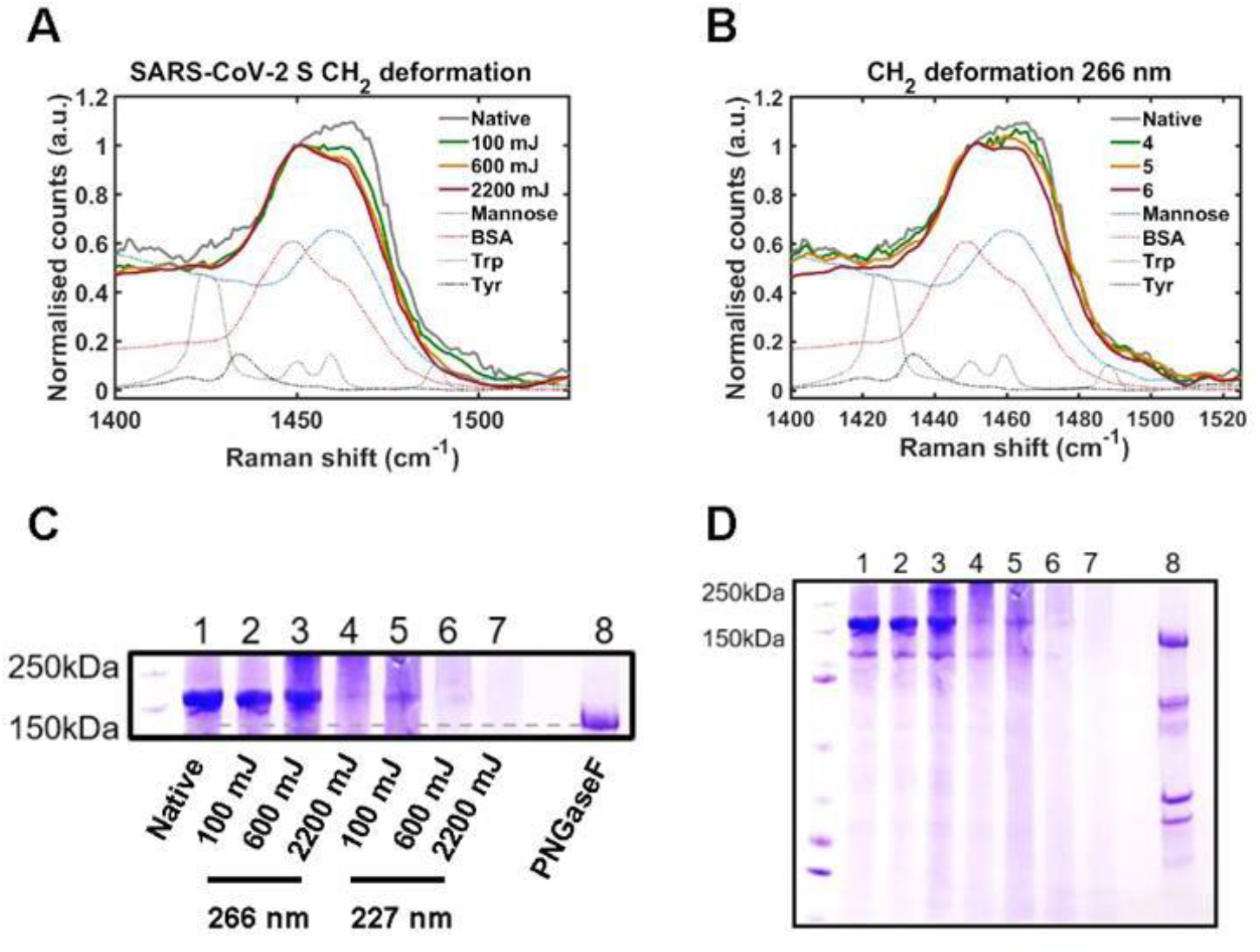
UVC irradiation of SARS-CoV-2 does not cause a decrease of glycosylation. A) Changes in glycan CH_2_ deformation induced by 227nm radiation and measured by Raman spectroscopy. B) Changes in glycan CH_2_ deformation induced by 266nm radiation and measured by Raman spectroscopy. n=2, plotted spectra represent the class means of 10-15 spectra per class. C) SDS PAGE analysis of SARS-CoV-2 glycosylation status. D) Full gel of data shown in C. n=1.

**Supplementary Figure 13.**
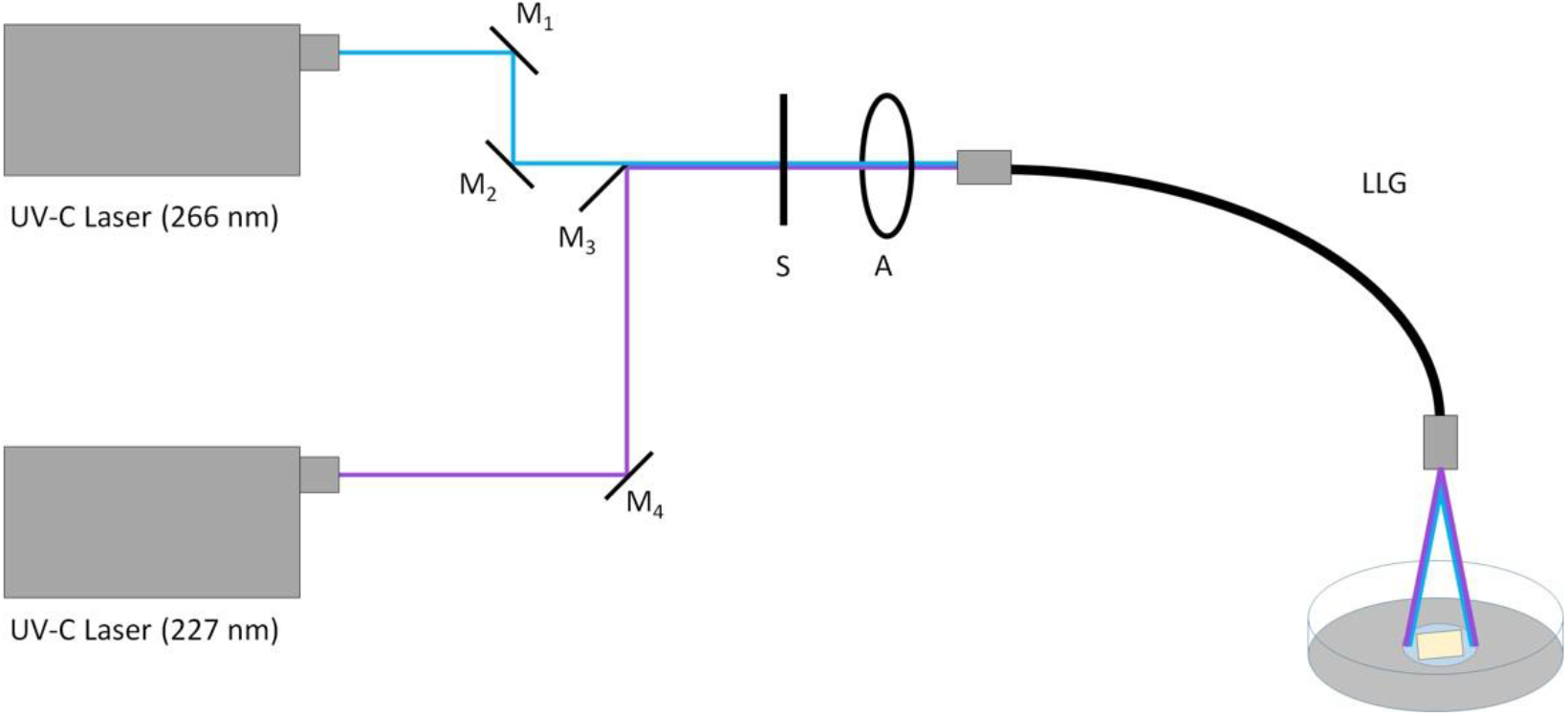
Diagram of dual-laser UV-illumination setup. Two UVC lasers, emitting at 227nm and 266nm, were coupled through an electronic shutter and adjustable aperture, and then into a liquid light guide (LLG). The large aperture of the LLG allowed the lasers to be coupled easily. Aluminium mirrors (M1-M4) were used to steer the laser beam. M3 was mounted on a translation stage to allow changing the UVC radiation wavelength incident on the samples.

**Supplementary Figure 14.**
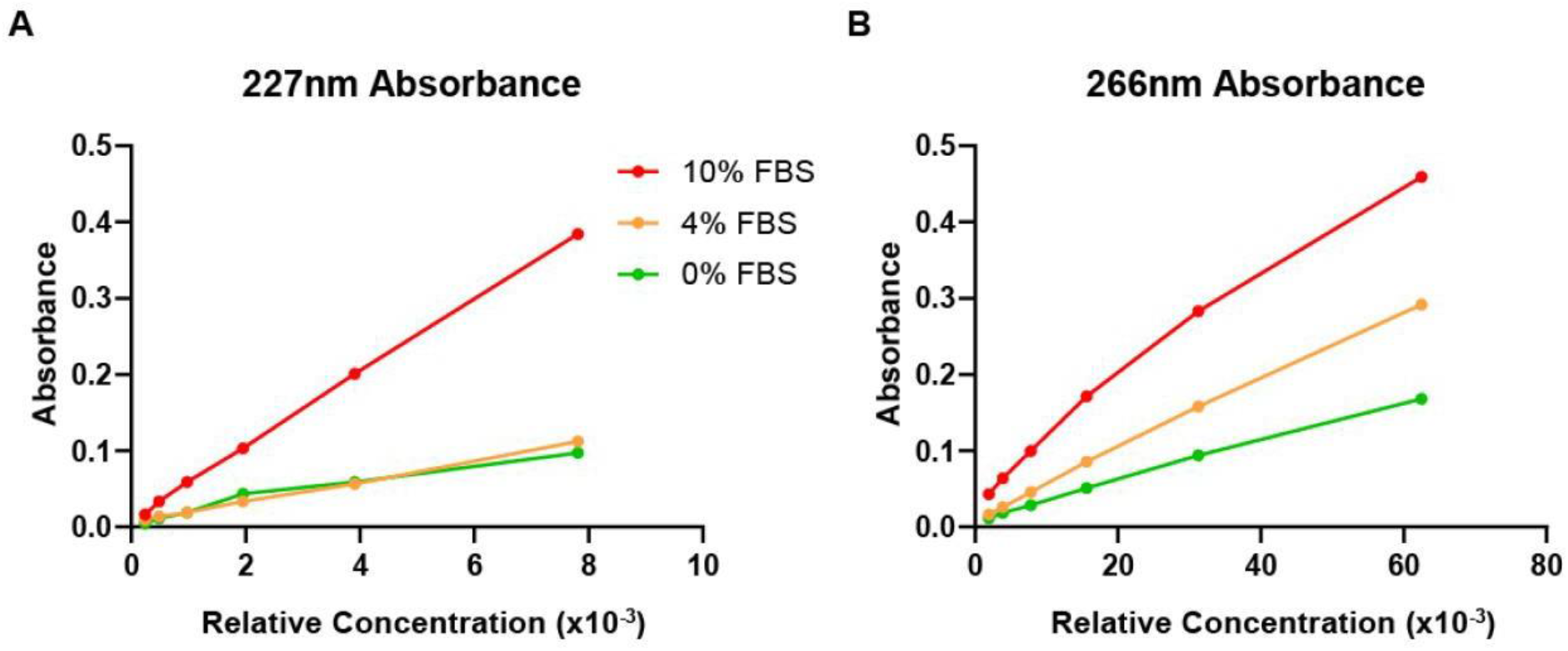
Absorbance of DMEM at 227nm and 266nm. Absorbance at 227nm and 266nm of DMEM supplemented with 0%, 4% and 10% Foetal Bovine Serum (FBS). Molar extinction coefficients (ε) calculated from this were used to calculate the effective dose received through a sample of a given thickness.

## Footnotes

a Operational constraints in the containment level 3 laboratory meant that altering radiant power within one experiment was not feasible.

